# Structural insights into the conformational plasticity of the full-length trimeric HIV-1 envelope glycoprotein precursor

**DOI:** 10.1101/288472

**Authors:** Shijian Zhang, Wei Li Wang, Shuobing Chen, Maolin Lu, Eden P. Go, Robert T. Steinbock, Haitao Ding, Heather Desaire, John C. Kappes, Joseph Sodroski, Youdong Mao

## Abstract

The human immunodeficiency virus (HIV-1) envelope glycoprotein (Env) trimer mediates viral entry into cells and is the major target for the host antibody response. In infected cells, the mature Env [(gp120/gp41)_3_] is produced by cleavage of a trimeric gp160 precursor. Proteolytic cleavage decreases Env conformational flexibility, allowing the mature Env to resist antibody binding to conserved elements. The conformational plasticity of the Env precursor skews the humoral immune response towards the elicitation of ineffectual antibodies, contributing to HIV-1 persistence in the infected host. The structural basis for the plasticity of the Env precursor remains elusive. Here we use cryo-electron microscopy to visualize two coexisting conformational states of the full-length Env precursor at nominal resolutions of 5.5 and 8.0 Å. The State-P2 conformation features a three-helix bundle of the gp41 heptad repeat region in the core, but has disordered membrane-interactive regions. State-P1 trimers lack the three-helix bundle and instead retain ordered transmembrane and membrane-proximal external regions embracing a central cavity. Our structural data shed light on the unusual plasticity of the Env precursor and provide new clues to Env immunogen discovery.

---

Human immunodeficiency virus type 1 (HIV-1), the etiologic agent of acquired immunodeficiency syndrome (AIDS), utilizes a metastable envelope glycoprotein (Env) trimer to engage host receptors and enter target cells^1^. During synthesis in the virus-producing cell, the Env precursor trimerizes and is glycosylated and cleaved into gp120 and gp41 subunits^2^. Env is the only virus-specific target accessible to neutralizing antibodies and has evolved a protective ‘glycan shield’ and surface variability^3, 4^. On the membrane of primary HIV-1, Env exists in a pre-triggered conformation (State 1) that resists the binding of commonly elicited antibodies^5, 6^. Binding to the receptor, CD4, on the target cell releases the restraints that maintain Env in State 1, allowing transitions through a default intermediate conformation (State 2) to the pre-hairpin intermediate (State 3).^5, 7^ In the more “open” State-3 Env, a trimeric coiled coil composed of the gp41 heptad repeat (HR1) region is formed and exposed, as is the gp120 binding site for the second receptor, either CCR5 or CXCR4.^8–10^ Binding to these chemokine receptors is thought to promote the insertion of the hydrophobic gp41 fusion peptide into the target cell membrane and the formation of a highly stable six-helix bundle, which mediates viral-cell membrane fusion^11–13^.

The conformational flexibility of Env required for HIV-1 entry has created challenges for structural studies. Nonetheless, structural information on individual HIV-1 Env subunits or subcomplexes^11, 12, 14, 15^, stabilized soluble gp140 trimers^3, 4, 16–20^, solubilized membrane Env trimers^21–23^ and virion Envs^24–26^, in either ligand-bound or free conformations, has been obtained by crystallography or electron microscopy. High-resolution structures of the HIV-1 Env ectodomain complexed with antibodies reveal a trimeric architecture stabilized by a gp41 HR1 coiled coil, from which emanates a three-bladed propeller composed of gp120 subunits^17, 23, 27^. Recent observations (Lu *et al*., unpublished observations) challenge the assumption that these trimer structures represent State 1, underscoring the need for additional studies relating Env structures to functional states on the virus entry pathway.

Env conformational plasticity may help HIV-1 avoid neutralization by antibodies^28, 29^. High titers of antibodies against State-2/3 Env conformations are elicited early during HIV-1 infection, but these antibodies cannot access their epitopes once the virus has bound CD4 and therefore do not neutralize efficiently^30, 31^. Antibodies that recognize conserved structures on State-1 Envs are generated less frequently and only after years of HIV-1 infection^32–34^. Antibodies with State-2/3 specificity often recognize the Env precursor more efficiently than cleaved Env^35–41^. Crosslinking the Env precursor exerted an effect on Env antigenicity similar to that of gp120-gp41 cleavage, suggesting that the Env precursor might be more flexible than mature Env.^29^ Here we use several approaches, including single-molecule FRET (smFRET) and single-particle cryo-electron microscopy (cryo-EM), to investigate the conformational landscape of a full-length, fully glycosylated Env precursor. Our analysis suggests that the Env precursor spontaneously samples multiple conformations related to States 1-3, marked by either the presence or absence of the central three-helix bundle of gp41. We discuss the implications of Env precursor conformational plasticity for HIV-1 evasion of the host immune response.

## Results

### Analysis of HIV-1 Env precursor conformations

Cleavage of the HIV-1 Env precursor affects its antigenicity.^35–41^ The recognition of the uncleaved and mature HIV-1_JR-FL_ Envs on the surface of transfected 293T cells exhibited distinct patterns for State 1-preferring broadly neutralizing antibodies (bNAbs) and State 2/3-preferring poorly neutralizing antibodies (Fig. 1a). Whereas the uncleaved Env bound antibodies capable of recognizing all three states, the mature Env bound only the potently neutralizing antibodies with State-1 preferences. The uncleaved Env apparently samples multiple conformations, but the mature Env exhibits a conformation that excludes the binding of weakly neutralizing antibodies.

**Fig. 1.**
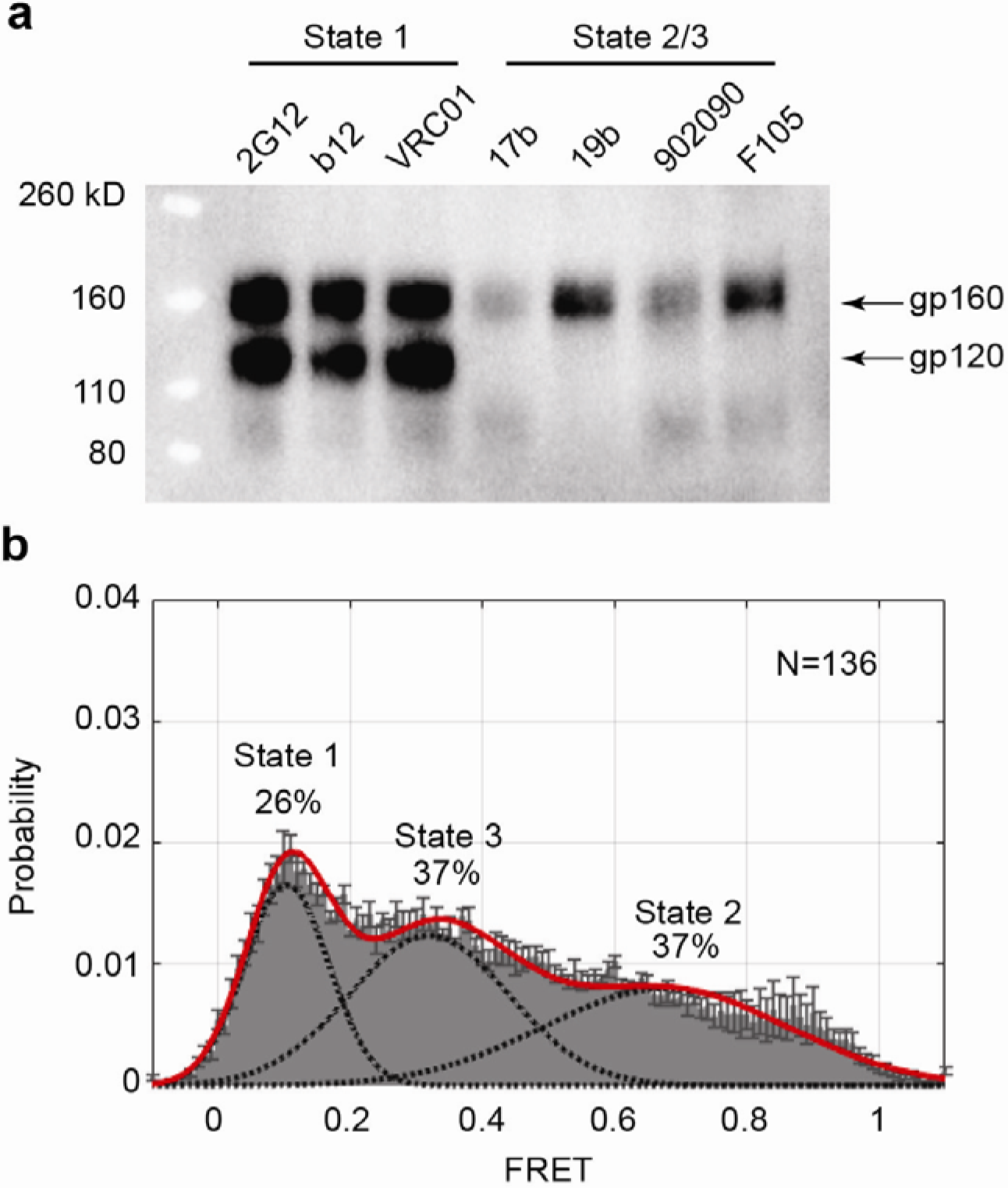
Conformational states of the HIV-1 Env precursor. **a**, HOS cells transiently expressing the wild-type HIV-1_JR-FL_ Env, which is incompletely cleaved in these cells, were incubated with the indicated State 1-preferring bNAbs or the State2/3-preferring weakly neutralizing antibodies. After washing and lysis of the cells, the antibody-Env complexes were purified using Protein A-Sepharose beads and analyzed by Western blotting with a rabbit anti-gp120 polyclonal serum. **b**, HIV-1_JR-FL_ Env(-) with V1 and V4 labels was purified from CHO cell membranes using a protocol identical to that used for preparation of Env(-) for cryo-EM imaging. The purified Env(-) was labeled and analyzed by smFRET. FRET trajectories were compiled into a population FRET histogram and fit to the Gaussian distributions associated with each conformational state, according to a hidden Markov model^5^. The percentage of the population that occupies each state as well as the number of molecules analyzed (N) is shown. The error bars represent the standard deviation from three independent data sets.

The above results are consistent with smFRET analyses of mature and cleavage-defective HIV-1_JR-FL_ Envs. The mature wild-type HIV-1_JR-FL_ Env predominantly samples a pre-triggered State-1 conformation; for example, the HIV-1_JR-FL_ Env on virions exhibited 53% State-1, 19% State-2 and 28% State-3 conformations^5, 7^. By contrast, the HIV-1_JR-FL_ Env(-) variant with an altered gp120-gp41 cleavage site samples all three conformations, exhibiting 23% State-1, 33% State-2 and 44% State-3 conformations (See Lu *et al*., accompanying manuscript). Thus, relative to the wild-type HIV-1_JR-FL_ Env, Env(-) exhibits an increased occupancy of the high-FRET (State-2) and intermediate-FRET (State-3) conformations.

The HIV-1 entry inhibitor, BMS-806, hinders transitions from State 1 and increases the occupancy of State 1 by the mature HIV-1 Env.^5, 35^ We recently showed that incubating BMS-806 with virions containing uncleaved Env(-) significantly enriched the low-FRET State-1 configuration, resulting in a conformational spectrum closer to that of the mature HIV-1_JR-FL_ Env; for example, BMS-806-treated HIV-1_JR-FL_ Env(-) on virions exhibited 58% State-1, 24% State-2 and 18% State-3 conformations (See Lu *et al*., accompanying manuscript). BMS-806 treatment of Env(-)-expressing cells resulted in significant decreases in recognition of the cell-surface Env by antibodies (e.g., b6, F105) exhibiting a preference for State 2, whereas the binding of bNAbs like 2G12 and VRC01 was less affected (Supplementary Fig. 1). After BMS-806 treatment and crosslinking with bis(sulfosuccinimidyl) suberate (BS3), cell-surface Env(-) was recognized efficiently by bNAbs (2G12, b12 and VRC01) but less well by poorly neutralizing antibodies. In contrast with the effects of BMS-806 treatment, incubation with dodecameric soluble CD4 has been shown to drive Env(-) on recombinant virions primarily into the intermediate-FRET configuration (State 3) (See Lu *et al*., accompanying manuscript). Thus, the occupancy of States 1 and 3 by Env(-) can be increased by the binding of BMS-806 or CD4, respectively, as was seen for the wild-type HIV-1_JR-FL_ Env.^5^

To gain additional understanding of the varied conformations sampled by the Env precursor, we purified full-length HIV-1_JR-FL_ Env(-) trimers from membranes of inducibly expressing CHO cells (Supplementary Fig. 2, a and b). The CHO cells were incubated with 10 µM BMS-806 during Env(-) synthesis; BMS-806 treatment of the Env(-)-expressing cells reduced the synthesis of Env(-) glycoforms enriched in complex carbohydrates, eliminating a presumably misfolded Env(-) subset displaying more surface-accessible glycans (Supplementary Fig. 2,c-e). Membranes containing Env(-) were incubated with saturating concentrations of BMS-806 and cross-linked with BS3 prior to solubilization by Cymal-5. The detergent in the protein solution was exchanged to a mixture of 4.5 mg/ml amphipol A8-35 and 0.005% Cymal-6 prior to cryo-EM imaging and data collection. Parallel smFRET studies estimated that 26% of detergent-solubilized Env(-) was in a low-FRET conformation consistent with State 1 and 37% in a high-FRET conformation resembling State 2; the remainder of the Env(-) molecules exhibited an intermediate-FRET (State-3-like) conformation (Fig. 1b).

### Cryo-EM structure determination

We collected cryo-EM data in video frames of a super-resolution counting mode with the Gatan K2 Summit direct electron detector mounted on an FEI Tecnai Arctica operating at 200 kV. We extracted 1,366,095 single-particle images of Env(-) from 10,299 drift-corrected movies using a deep-learning-based approach in a template-free fashion^42^ (Fig. 2a). These particle images were subjected to exhaustive unsupervised 2D classification by a statistical manifold learning algorithm^43^ and unsupervised maximum likelihood-based 3D classification^44^ (Supplementary Figs. 3 and 4). These resulted in several 3D classes, which can be grouped into two distinct major architectures: State P2 with a gp41 three-helix bundle (3-HB_c_) in the trimer center and State P1 lacking the central 3-HB_c_ and instead exhibiting a void surrounding the trimer axis. State P2 trimers generally exhibited slightly smaller sizes than State P1 trimers, and were more abundant in the Env(-) population (∼89% State P2-like versus ∼11% State P1-like; see Supplementary Fig. 3). We chose the most homogeneous 3D class in each state and refined the maps of States P1 and P2 to 8.0 and 5.5 Å, respectively (Supplementary Fig. 4). We did not observe density associated with the ∼140-residue Env cytoplasmic tail in any of the 3D classes, indicating that this element is largely disordered in these preparations.

**Fig. 2.**
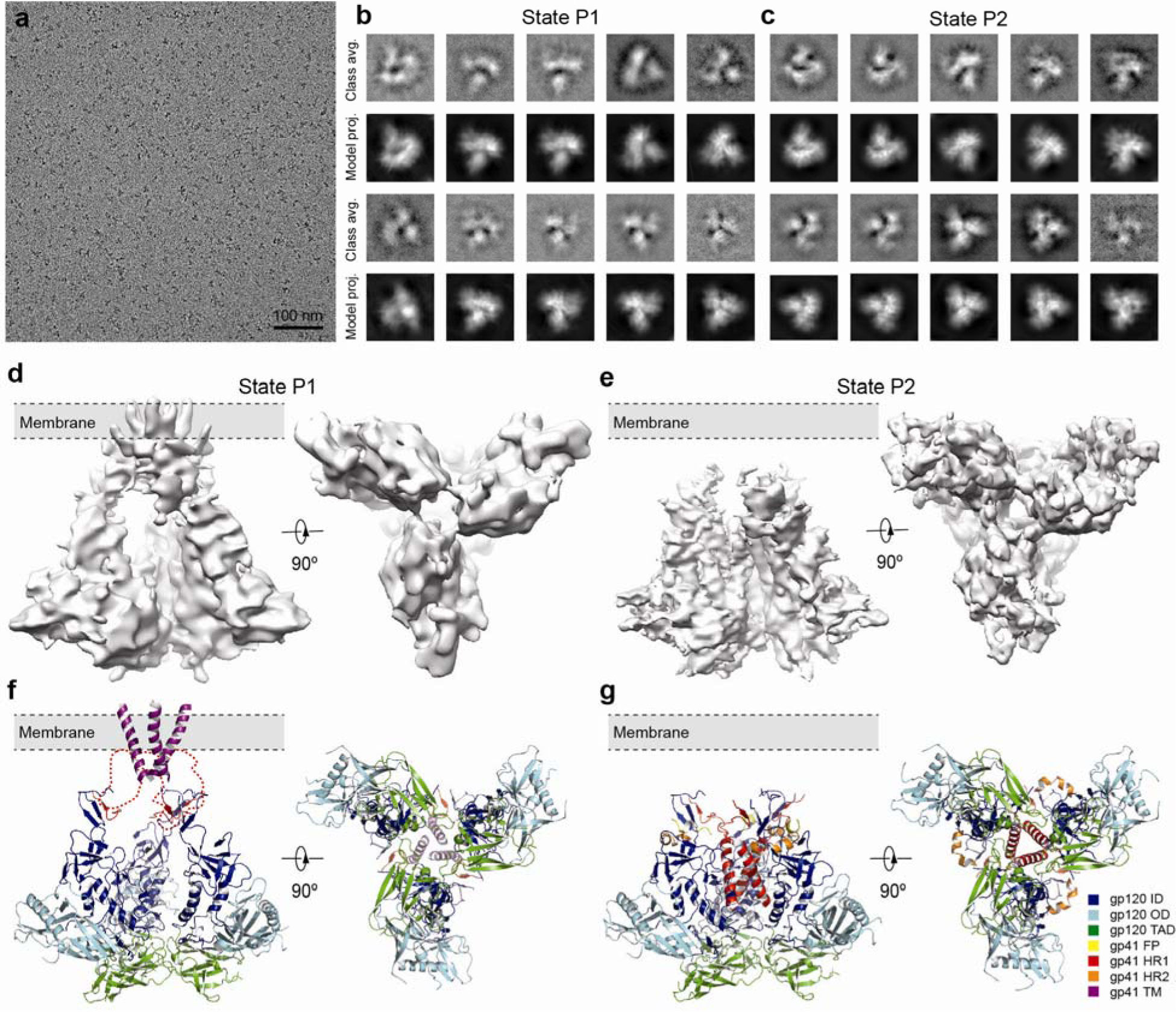
Cryo-EM analysis of the full-length HIV-1_JR-FL_ Env(-) trimer in two alternative conformations. **a**, A typical cryo-EM micrograph of Env(-) trimers taken with the Gatan K2 direct electron detector. **b**, Unsupervised State-P1 class averages calculated by ROME software in comparison with their corresponding model projections**. c**, Unsupervised State-P2 class averages calculated by ROME software in comparison with their corresponding model projections**. d**, Cryo-EM map of State P1 in two orthogonal views. **e**, Cryo-EM map of State P2 in two orthogonal views. **f**, The pseudo-atomic model of State P1 in two orthogonal views. The dashed red line marks the FP/HR1/HR2/MPER regions of the gp41 subunit that were not modeled. **g**, The pseudo-atomic model of State P2 in two orthogonal views. Color codes: ID, inner domain, blue; OD, outer domain, cyan; TAD, trimer association domain, green; FP, fusion peptide, yellow; HR1, heptad repeat 1, red; HR2, heptad repeat 2, orange; TM, transmembrane region, purple.

**Fig. 3.**
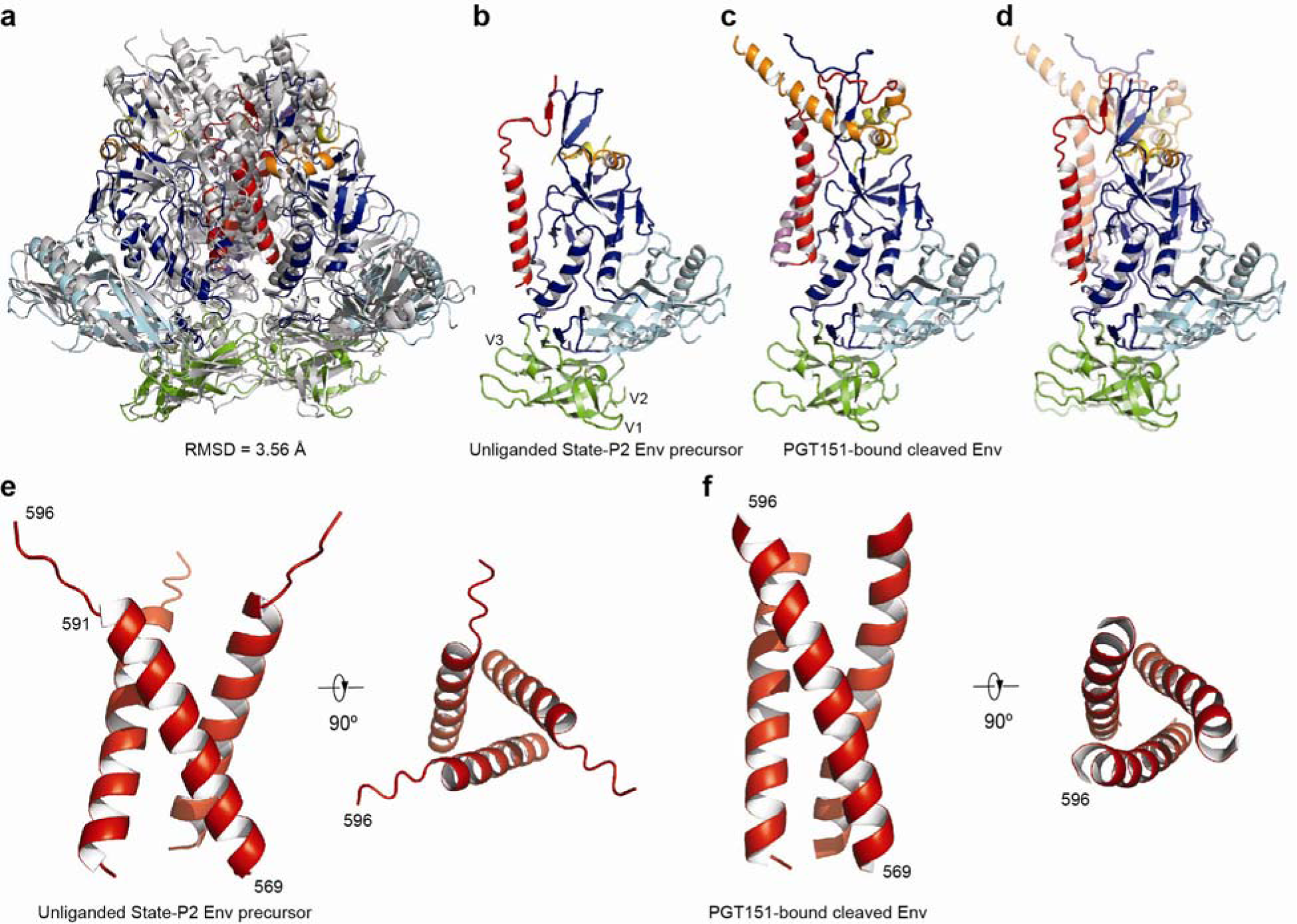
Comparison of the structures of the HIV-1_JR-FL_ State-P2 Env(-) and PGT151-bound Env(+)ΔCT trimers. **a**, The pseudo-atomic model of State-P2 Env(-), colored by domain code as in Figure 2, is superposed with that of the PGT151-bound Env(+)ΔCT, colored grey. **b**, Side view of the pseudo-atomic model of the gp160 protomer of the Stat Env(-) trimer. **c**, The same view of the pseudo-atomic model of the gp120/gp41 protomer of the PGT151-bound Env(+)ΔCT. **d**, The protomer structures of State-P2 Env(-) and Env(+)ΔCT are superimposed, after alignment of the gp120 outer domain. The structure of the Env(+)ΔCT protomer is depicted with 50% transparency to facilitate differentiation from the structure of State-P2 Env(-) protomer. **e**, The central 3-HB_c_ formed from part of HR1 in the State-P2 Env(-) trimer in two orthogonal views. **f**, The central 3-HB_c_ formed from part of HR1 in the PGT151-bound Env(+)ΔCT trimer in two orthogonal views.

**Fig. 4.**
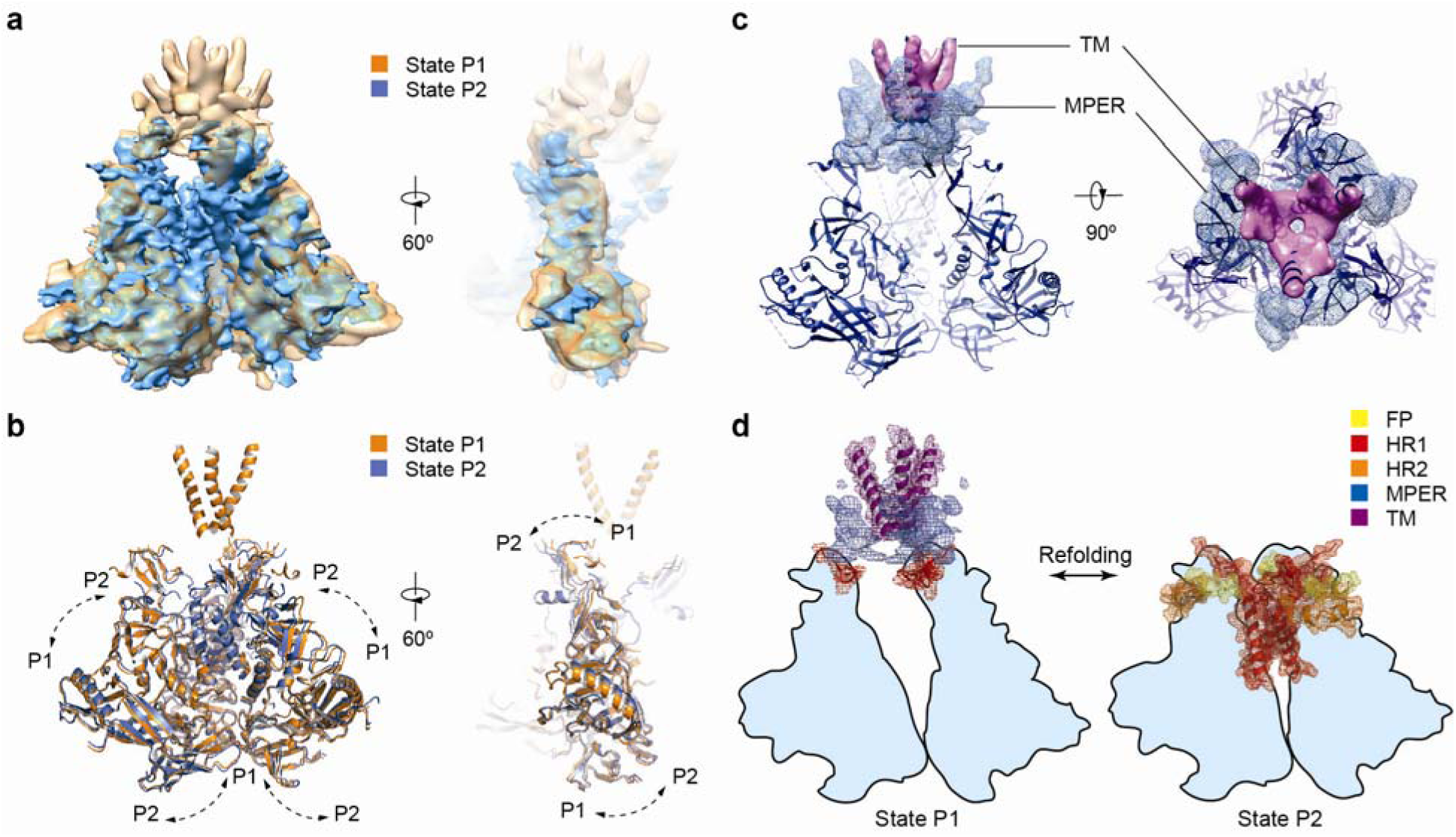
Comparison of the Env(-) structures in States P1 and P2. **a**, The cryo-EM maps of the Env(-) States P1 and P2 are superposed and viewed from two different perspectives. **b**, The pseudo-atomic models of the Env(-) States P1 and P2 are superposed and viewed from two different perspectives, showing that the gp120 subunits are rotated with respect to each other. **c**, The segmented densities associated with the gp41 transmembrane (TM) region (magenta solid surface) and the membrane-proximal external region (MPER) (blue mesh) in State P1 are displayed. The pseudo-atomic model of the State-P1 Env(-) trimer (deep blue ribbon) is superposed. **d**, The model illustrates the refolding of Env(-) between States P1 and P2. Except for the MPER density segmentation (blue mesh), the rest of the gp41 elements are drawn with their mesh representation of the pseudo-atomic models of P1 and P2.

Figures 2b and c show the typical reference-free CTF-corrected class averages of Env(-) in States P1 and P2, respectively, compared with their corresponding model projections. The side views of State P1 and P2 emphasize the fundamental difference in their architecture (Fig. 2d and e). Our State-P2 map generated by the unsupervised maximum-likelihood-based 3D classification exhibited high-resolution structural features agreeing with the crystal structures of sgp140 SOSIP.664,^3, 4, 17, 27, 45^ providing direct validation of our unsupervised 3D classification. Although the State-P1 map differs from the State-P2 map, it was achieved by the same unsupervised 3D classification procedure. Starting with the high-resolution crystal structures of the sgp140 SOSIP.664 Env trimer, we fit a pseudo-atomic model for the 5.5-Å map of State P2 (Fig. 2g, Supplementary Table 1). The pseudo-atomic model of the gp120 subunit from State P2 was then used as a starting model to derive a pseudo-atomic model for State P1 (Fig. 2f, Supplementary Fig. 4).

To confirm that the unsupervised 3D classification did not overfit the data, we conducted additional validation tests. First, a tilt-pair validation with data collected with the K2 direct electron detector was performed by incorporating a 3D classification procedure into the tilt-pair analysis (see Methods). The results validated the P1 map resulting from our unsupervised 3D classification (Supplementary Fig. 5). Second, to exclude potential reference bias in cryo-EM refinement, we conducted high-resolution refinement of the P1 dataset using the P2 map as an initial model, and vice versa. The P1 map refined using the P2 initial model is virtually identical to the P1 map refined using its own low-passed filtered model (Supplementary Fig. 6). Likewise, the P2 map refined using the P1 initial model conforms to the P2 map derived by refinement using a consensus model. These validation tests support the conclusion that the P1 and P2 maps represent distinct conformations existing in the purified Env(-) sample.

### Overview of Env precursor conformational states

The structure of State-P1 Env(-) shares a topology similar to that of the cryo-electron tomography (cryo-ET) map of the native Env spike on HIV-1 virions^25^ and closely reproduces the molecular surface of the uncleaved Env with the cytoplasmic tail (CT) deleted (Env(-)ΔCT) (Supplementary Fig. 7).^22^ The central cavity surrounding the trimer axis is a prominent feature of these structures. Peripheral to the transmembrane helices, prominent tripod-like structures project toward the membrane in State P1. The three blades of gp120 in State P1 are slightly more splayed outward than in State P2 (Fig. 2d-g).

The structure of State-P2 Env(-) shares a similar topology to that of sgp140 SOSIP.664 (refs. 3, 4, 17, 18, 27, 45) and the cleaved Env(+)ΔCT, the latter structure determined in complex with the PGT151 broadly neutralizing antibody (bNAb).^23^ A 3-helix bundle (3-HB_c_) formed by the C-terminal portion of the gp41 HR1 region is a central feature of these structures (Fig. 3). The gp120 component of the sgp140 SOSIP.664 and Env(+)ΔCT-PGT151 trimers has been shown to assume a State-2 conformation (Lu *et al*., unpublished observations). Thus, the compatibility of our State-P2 map with these structures suggests that the global conformation of the State-P2 Env precursor resembles State 2 of the mature Env.

### Structure of the Env precursor in State P2

Despite the overall similarity between our State-P2 HIV-1_JR-FL_ Env(-) structure and the PGT151-bound cleaved HIV-1_JR-FL_ Env(+)ΔCT structure, the two conformations exhibit some notable differences (Fig. 3). First, the State-P2 Env(-) gp120 subunits are shifted away from the trimer axis relative to their position in the PGT151-bound Env(+)ΔCT structure (Fig. 3a). When the gp120 structures are aligned using the gp120 outer domain (OD), the gp120 inner domain (ID) exhibits a counterclockwise rotation in State-P2 Env(-) relative to the PGT151-bound Env(+)ΔCT (Fig. 3b-d).

Second, the gp120 trimer association domain (TAD) adopts a flatter conformation in the State-P2 Env(-). The gp120 TAD, composed of the V1, V2 and V3 regions, is situated at the trimer apex and can influence the propensity of Env to move from State 1.^7^ In the State-P2 structure, the V1/V2 region assumes a four-stranded Greek-key β-sheet topology, as seen in the crystal structures of sgp140 SOSIP.664 and V1/V2 scaffolds complexed with bNAbs PG9 and PG16 ^19, 46^. Unlike the vertical configuration in the PGT151-bound Env(+)ΔCT, the inter-V1V2 loop (residues Ser164 to Glu168) mediating gp120 inter-subunit interaction at the trimer apex^17, 27, 47^ adopts a flatter and tighter configuration in State P2, stacking over the V3 tip. Consequently, the V1 loop of State-P2 Env(-) slides outwards away from trimer axis by about 5 Å. Thus, the quaternary structure of the gp120 TAD is overall more splayed out at the apex of the State-P2 Env(-).

Third, the central 3-HB_c_ formed by part of gp41 HR1 is about 5 residues shorter in the State-P2 Env(-) than in the PGT151-bound Env(+)ΔCT (Fig. 3e,f). The crossing angle of the three helices is also slightly larger in the State-P2 structure. Coincidently, the central 3-HB_c_ is about 6 Å closer to the gp120 TAD apex in the State-P2 Env(-) than in the PGT151-bound Env(+)ΔCT. Thus, the central 3-HB_c_ in the PGT151-bound Env(+)ΔCT trimer is tighter and apparently more stable than that in the State-P2 Env(-). These observations are compatible with the movement of gp120 away from the trimer axis in the State-P2 Env(-), relative to its position in the PGT151-bound Env(+)ΔCT trimer.

Fourth, major differences were also found in the N-terminal portion of the HR1 region and the entire HR2 region of gp41. Similar to the sgp140 SOSIP.664 Env structure, the N-terminal portion of HR1 is not resolved in the State-P2 Env(-) map. However, this region is helical in the PGT151-bound Env(+)ΔCT structure, allowing HR1 to adopt a helix-turn-helix topology^23^. Notably, the long gp41 HR2 helices observed in the sgp140 SOSIP.664 and PGT151-bound Env(+)ΔCT structures are not evident in the State-P2 Env(-) map.

The gp41 membrane-proximal external region (MPER) and transmembrane (TM) regions were truncated in the sgp140 SOSIP.664 Env construct, and were also found to be disordered in the PGT151-bound Env(+)ΔCT structure. Consistent with this observation, in our State-P2 Env(-) map, these regions were not well resolved.

### Structure of the Env precursor in State P1

The architecture of the State-P1 Env precursor is notably different from that of State P2 in two respects. First, the striking hollow architecture of the State-P1 Env(-) trimer, with a central cavity along the trimer axis, contrasts with the more compact State-P2 architecture. Second, the membrane-proximal regions of State-P1 Env(-) are mostly ordered, in contrast to the disorder of these regions in State P2. Although the resolution of the P1 map did not permit an accurate Cα backbone trace, the pseudo-atomic model of the gp120 subunit derived from the P2 map could be fitted into the P1 map as a rigid body. The NMR structure of a single gp41 membrane-spanning helix^15^ was fit into the transmembrane region of the P1 map with flexible adjustment. In the final pseudo-atomic model of State P1, the majority of the gp41 ectodomain was not interpreted due to lack of resolution.

The orientation of the gp120 subunits differs slightly in the State-P1 and State-P2 Env(-) trimers. In the sgp140 SOSIP.664 and State-P2 structures, the V3 loop, forming an antiparallel β-hairpin situated beneath the V1/V2 elements, contributes to the inter-protomer interaction among gp120 subunits^17, 27, 47^. The interaction of the tip of the V3 β-hairpin with the neighboring gp120 subunit forms a pivot that allows each gp120 core to adjust its tilt along the trimer axis in different conformational states. Using the V3 β-hairpin as a pivot, each gp120 subunit is rotated outward, away from the trimer axis, in State P1 relative to State P2 (Fig. 4a, b). When viewing the two conformations laterally, the entire gp120 blade in State P1 is rotated in a direction that is better aligned with the trimer axis (Fig. 4b, right panel). Such structural rearrangement in the Env(-) trimer is facilitated via the gp120 inner domain, which exhibits a layered architecture that has been shown to assist CD4 binding by promoting Env conformational transitions from State 1 to States 2/3.^48^

In contrast to State P2, the three transmembrane helices in the State P1 Env(-) trimer are well ordered and form a left-handed coiled coil, in line with the cryo-EM structure of an unliganded Env(-)ΔCT trimer^22^ and the recent NMR structure of a trimeric gp41 transmembrane peptide^15^. The State P1 Env(-) transmembrane coiled coil seems more splayed out on the cytoplasmic side of the membrane as compared to that in the Env(-)ΔCT trimer^22^ or in the NMR structure of the gp41 transmembrane trimer^15^, possibly related to the disorder in the Env(-) cytoplasmic tails. A density extending from the transmembrane helices towards the cytoplasmic side of the membrane may indicate a mobile, intramembrane charged residue, arginine 596. Consistent with the cryo-EM reconstruction of Env(-)ΔCT, the transmembrane region is encircled with triangular layers of MPER structures that potentially abut the membrane^22^ (Fig. 4c). In the P1 map, the HR2 region appears to be broken into several short secondary structure elements that interdigitate with the MPER and likely with HR1 as well^22^.

### Transitions between State P1 and State P2

Reversible transitions between State P1 and State P2 of the HIV-1 Env precursor occur spontaneously and in response to particular ligands (See Lu *et al*., accompanying manuscript). The sharp contrast between the ordered gp41 regions of State P1 and State P2 suggests that the gp41 ectodomain can undergo refolding around its central regions, a process that dictates a tight association between a specific gp41 quaternary structure and a given state (Fig. 4d). Such gp41 refolding involves movement of the gp41 HR1 C-terminal portion from more peripheral locations remote from the trimer center in State P1 to the 3-HB_c_ surrounding the trimer axis in State P2. In State P1, concomitant with more peripherally located HR1 elements and a more stable quaternary architecture of TM/MPER/HR2, the three gp120 subunits fall slightly apart with an inter-protomer distance greater than that in State P2 (Fig. 4a,b). As the HR1 refolds into the 3-HB_c_ during the transition from State P1 to State P2, the 3-HB_c_ structure pulls each gp120 closer to the trimer axis.

Movement of gp120 may be assisted by particular structures. In addition to the rigid body motion of the gp120 outer domain described above, conformational changes likely involve the gp120 trimer association domain (TAD) and inner domain^5, 48^. The low resolution of the State-P1 Env(-) structure precludes a definitive comparison of these structural regions in P1 and P2 states. However, flanking the central cavity of State P1 is a prominent loop-like structure that may represent part of the gp120 inner domain.^48^ If this hypothesis is correct, movement of gp120 is necessitated by 3-HB_c_ formation, which in State 2 occupies the same space as the loop-like structure. Movement of gp120 coincides with a break in the quaternary structure of the membrane-proximal gp41 regions (TM/MPER/HR2) (Fig. 4d). Although the extent of the latter changes in gp41 will likely be influenced by the presence of the membrane, the results suggest a relationship between state transitions in the rest of the Env precursor and the conformation of the membrane-proximal elements of gp41.

### Env precursor glycosylation

Given that State-P1 and State-P2 Env(-) trimers readily interconvert, we expect their glycosylation profile to be similar; however, properties of the Env glycan shield could differ as a result of Env conformation-dependent shifts in the location of particular glycans. In the State-P2 map, we identified the majority of the gp120 glycan-associated densities, including some of those identified as complex glycans by mass spectrometric analysis^49^ of the purified Env(-) glycoprotein (Supplementary Fig. 2e). Consistent with the crystallographic analysis of the glycan shield in the sgp140 SOSIP.664 context^3, 4^, the gp120 glycans in high-density areas exhibit at least partial order in the peptide-proximal glycan residues. Most distal glycan residues are not well resolved, reflecting their dynamic nature and heterogeneity.

After pseudo-atomic interpretation of the gp120 components in the State-P1 map, there are several extra densities that are of the appropriate size and shape to be glycans. Most of these are positioned similarly to the State-P2 glycans; however, more specific characterization of the State-P1 glycan structure will require additional data. Likewise, although glycan-like densities are apparent in the gp41 subunit of State P1, interpretation awaits a more complete gp41 atomic model for this state.

## Discussion

This study highlights the conformational plasticity of the HIV-1 Env precursor and provides insights into its structural basis. Antibody or ligand binding and smFRET analyses indicate that the Env precursor can sample multiple conformations that resemble States 1, 2 and 3 of the mature viral Env spike. The conformational plasticity of the Env precursor contrasts with the behavior of the mature Env, which in the absence of ligands largely resides in State 1 (Fig. 6). Therefore, proteolytic cleavage stabilizes State-1 Env, which is highly resistant to neutralization by antibodies recognizing other Env conformations. Although proteolytic maturation also primes the membrane-fusing potential of other Class I viral membrane fusion proteins, the effects of cleavage on HIV-1 Env conformational plasticity are unusual. For example, crystal structures comparing the influenza virus precursor, HA0, with the cleaved HA1/HA2 trimer showed differences only in the immediate vicinity of the cleavage site^50^. Inefficient Env folding and cleavage in HIV-1-infected cells, which undergo extensive lysis from viral cytopathic effects and cytotoxic immune responses, would result in the abundant presentation of Env precursor antigens in conformations other than State 1. The resulting diversion of host antibody responses away from State-1 Env, the major target for neutralizing antibodies, would have considerable advantages for a persistent virus like HIV-1.

**Fig. 5.**
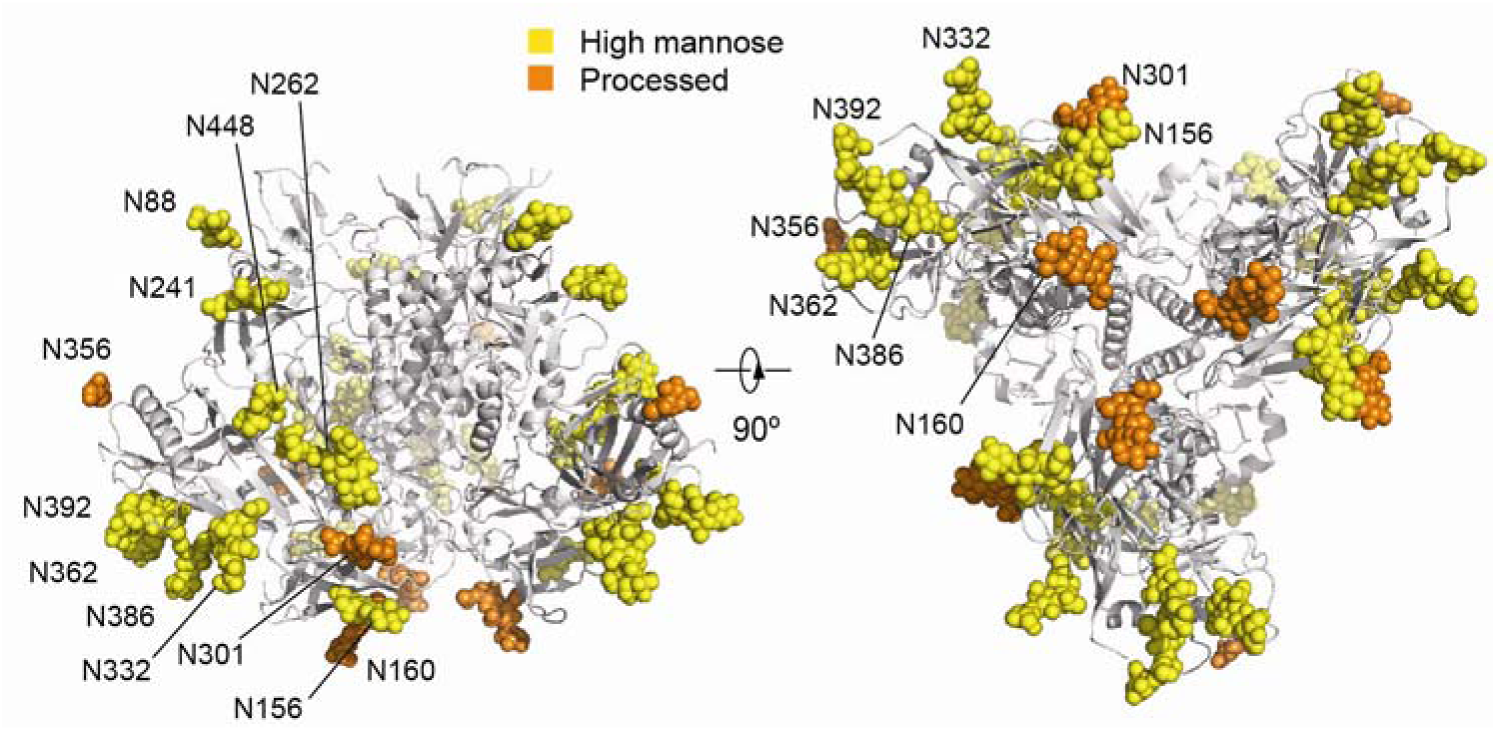
Env(-) glycosylation. The pseudo-atomic model of glycans in State-P2 Env(-) in two orthogonal views. The high-mannose glycans are colored yellow. The complex glycans are colored orange.

**Fig. 6.**
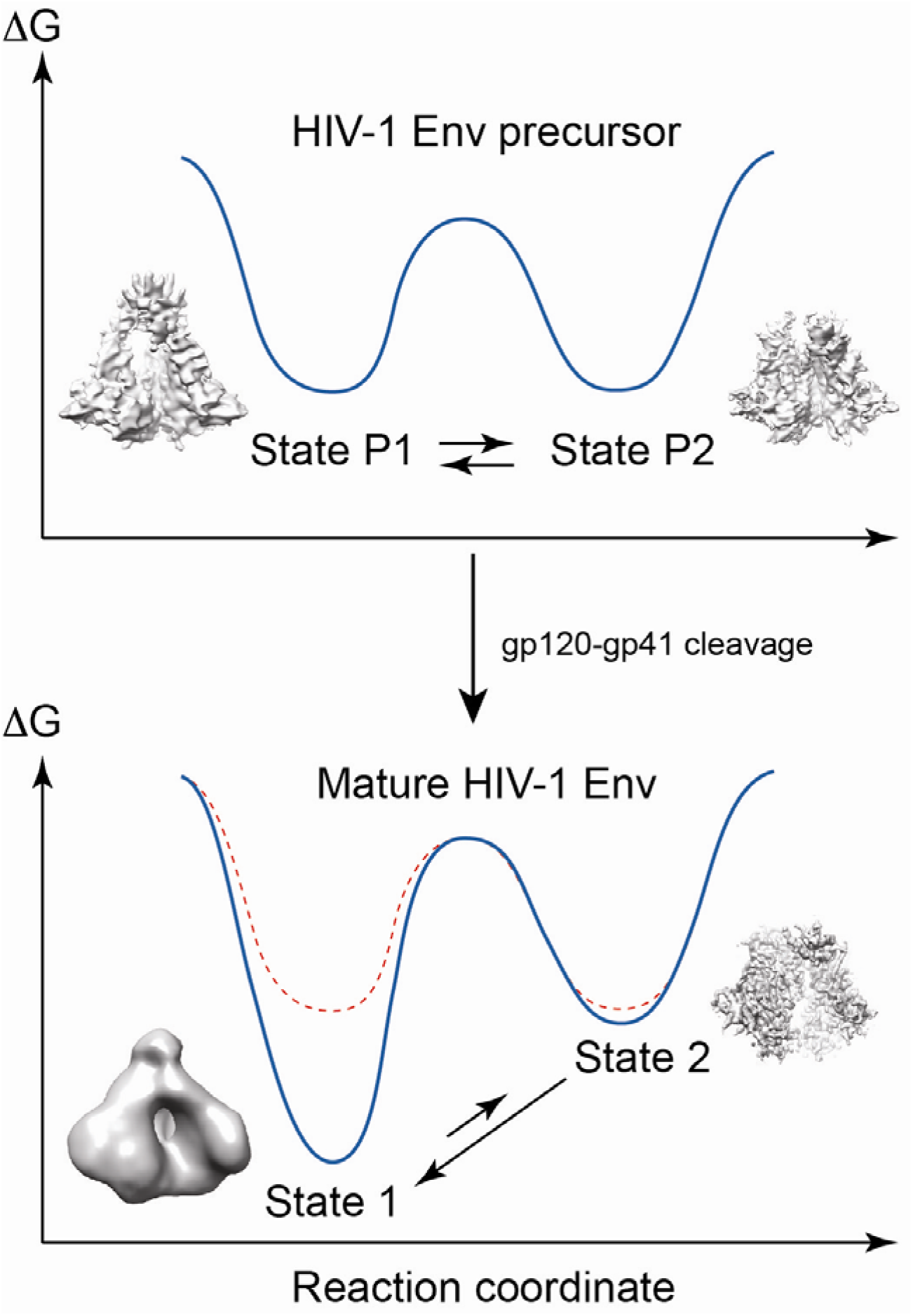
Proposed mechanistic model of changes in the HIV-1 Env energy landscape upon gp120-gp41 cleavage. Based on the occupancies of Env conformations measured by smFRET and assuming that the HIV-1 Env conformations are in equilibrium, the relative free energies of the uncleaved Env States P1 and P2 (upper panel) and the cleaved Env States 1 and 2 (lower panel) are depicted. The conformational plasticity of the HIV-1 Env precursor allows State P2 to be presented to the host immune system, skewing antibody responses away from State 1-like conformations. Env cleavage into gp120 and gp41 stabilizes State 1, allowing this conformation to predominate on the mature virion Env.

We identified experimental conditions that allowed us to visualize two distinct coexisting conformations of the uncleaved Env(-) trimers. The entry inhibitor BMS-806, which blocks Env conformational transitions,^5, 35^ was used to enrich State P1 in the preparation. The visualization of two coexisting states within a single mixture of Env(-) molecules, likely corresponding to two of the three documented Env precursor states, substantiates the diversity of Env precursor conformations suggested by smFRET and antigenicity studies. Only the pre-hairpin intermediate conformation (State P3) normally induced by the binding of multiple CD4 molecules was not visualized.

Although further work is needed to understand the effects of gp120-gp41 cleavage on the conformations of the membrane Env trimer, the P1 and P2 Env precursor states observed here are likely related to the mature State-1 and State-2 Envs, respectively. The higher resolution of the State P2 Env(-) map allowed us to establish the structural similarity to the gp120 subunits and gp41 3-HB_c_ in the sgp140 SOSIP.664 and PGT151-Env(+)ΔCT trimers, which are proteolytically mature. The gp120 subunits of these trimers have recently been shown by smFRET analysis to assume a State 2-like conformation (Lu *et al*., unpublished observations). The repeated appearance of State-2/State-P2 gp120 structures, despite the different circumstances of Env trimer preparation, is consistent with the conclusion from virological studies that State 2 is a default conformation assumed when State 1 is destabilized^7^. Although the State-P2 Env(-), sgp140 SOSIP.664 and PGT151-Env(+)ΔCT structures all have a gp41 3-HB_c_, this coiled coil is shorter in the State-P2 Env(-) trimer. Other differences exist between the gp41 ectodomains of State-P2 Env(-) and the two other trimers, particularly in the HR2 region. Additional studies are required to explain the observed differences.

There are reasons to support the correspondence of the State-P1 Env(-) structure with that of the precursor of the mature State-1 Env. First, smFRET verified the existence of a State 1-like Env conformer in the BMS-806-treated Env(-) preparation. Second and most importantly, the State-P1 Env(-) structure resembles the structures of mature Env trimer spikes on HIV-1 virions. The HIV-1 Env spike, which was first observed through *in situ* cryo-ET on native virions^51^, displayed a tripod architecture in the membrane-spanning anchor region that was challenged and debated^26, 52–54^. Here, we visualized membrane-proximal tripod-like gp41 structures in State P1 that potentially buttress the Env spike. The observed variation in the splay and position of these tripod-like structures^26, 52–54^ indicates their conformational flexibility, explaining how they could be highly dependent on Env preparation variables and thus escape detection. Indeed, the entire gp41 MPER and TM regions were disordered in a high-resolution cryo-EM reconstruction of the PGT151-bound Env(+)ΔCT trimer^23^.

An independent cryo-ET study of mature Env on HIV-1 virions at an improved resolution, in contrast to the early cryo-ET study, revealed a prominent central cavity in the Env spike^25^. This architecture is consistent with our State-P1 conformation as well as our previous map of Env(-)ΔCT^22^. However, later single-particle cryo-EM studies of sgp140 SOSIP.664 trimers, which we now know represent a State-2 gp120 conformation (Lu *et al*., in preparation), failed to observe the central cavity, leading to its speculative reinterpretation as an “artefact” resulting from low resolution^16, 18^. Inconsistent with this interpretation, low-pass filtering our State-P2 map to the resolution of the cryo-ET study does not give rise to a prominent central cavity, in stark contrast to the central cavity consistently shown in the State-P1 map filtered at multiple different resolutions (Supplementary Fig. 8). Thus, we speculate that the cryo-ET model of the mature Env spike on native virions^25^ may represent an averaged State-1 conformation distinct from State-P2 and State-2 conformations. In summary, despite differences in approaches and experimental details, both mature and uncleaved HIV-1 Env trimers with a similar hollow architecture have been visualized repeatedly^21, 22, 25, 54^. The central cavity is suggestive of a high potential energy that is expected of the metastable State-1 Env and its immediate precursor.

The ability of HIV-1 Env to convert readily between the State-P1 and State-P2 conformations is essential to the success of Env precursor plasticity as a strategy for immune evasion. State P2-like conformations need to be adequately populated to serve as decoys to the host immune system, but State P1-like conformations are required to supply virions with mature functional Envs, which are almost exclusively in State 1 and therefore resistant to neutralization by most antibodies. Lack of gp120-gp41 cleavage must prevent intra-spike interactions that stabilize and maintain State 1; having the Env precursor and mature Env share a similar global architecture would allow rapid formation of optimal State 1-restraining bonds upon cleavage. On the other hand, State-P2 Env precursors must be able to revert to State P1 easily. As this process involves refolding the gp41 HR1 coiled coil, it may be facilitated by the shorter, less stable 3-HB_c_ in the State-P2 Env precursor.

HIV-1 Env trimers are metastable and potentially sample many conformations, some of which retain function. Previous investigations revealed different facets of this vast Env conformational space. Whether due to lack of exhaustive 3D classification^21, 22, 24–26, 52, 54^ or the use of specific ligands, trimer-stabilizing changes or preparation variables^4, 17, 18, 23, 27, 45^, these approaches led to the observation of specific Env conformations. More recent appreciation of HIV-1 Env dynamics has revealed new intermediates on the virus entry pathway, allowing assignment of some of these structures to functional or off-pathway states^7^. Further investigation is needed to fill gaps, particularly in our understanding of State-1 Env trimers, and to achieve a more complete picture of the functional Env conformational landscape.

## Methods

### Protein expression and purification

For expression of the uncleaved full-length membrane-anchored HIV-1_JR-FL_ Env(-) glycoprotein, the *env* cDNA was codon-optimized and was cloned into an HIV-1-based lentiviral vector. These Env sequences contain a heterologous signal sequence from CD5 in place of that of wild-type HIV-1 Env. The proteolytic cleavage site between gp120 and gp41 was altered, substituting two serine residues for Arg 508 and Arg 511. In the HIV-1_JR-FL_ Env(-) glycoprotein, the amino acid sequence LVPRGS-(His)_6_ was added to the C-terminus of the cytoplasmic tail. For Env(-) expression, the *env* coding sequences were cloned immediately downstream of the tetracycline (Tet)-responsive element (TRE). Our expression strategy further incorporated an internal ribosomal entry site (IRES) and a contiguous puromycin (puro) T2A enhanced green fluorescent protein (EGFP) open reading frame downstream of *env* (TRE-*env*-IRES-puro.T2A.EGFP). Uncleaved membrane-anchored Env(-) was produced by exogenous expression in CHO cells. Briefly, a vector encoding HIV-1_JR-FL_ Env(-)was packaged, pseudotyped with vesicular stomatitis virus (VSV) G protein, and used to transduce CHO cells (Invitrogen) constitutively expressing the reverse Tet transactivator (rtTA). Transduced cells were incubated for 24 h in culture medium containing 1 μg/ml of doxycycline (DOX) and were then selected for 5 to 7 days in medium supplemented with puromycin (10 μg/ml). High-producer clonal cell lines were derived using a FACSAria cell sorter (BD Biosciences) to isolate individual cells expressing high levels of EGFP. The integrity of the recombinant *env* sequence in the clonal lines was confirmed by sequence analysis of PCR amplicons. Clonal cultures were adapted for growth in a serum-free suspension culture medium (CDM4CHO; Thermo Fisher).

For the exogenous production of the Env(-)glycoprotein, cells were expanded in a suspension culture using a roller bottle system (Thermo) and were treated with 1 μg/ml of DOX and 10 μM BMS-378806 (herein referred to as BMS-806) (Selleckchem) after reaching a density of >4 × 10^6^ cells per ml. After 18 to 24 h of culture with DOX and BMS-806, the cells were harvested by centrifugation. During the remainder of the purification procedure, 10 μM BMS-806 was added to all buffers. The cell pellets were homogenized in a homogenization buffer (250 mM sucrose, 10 mM Tris-HCl [pH 7.4], and a cocktail of protease inhibitors [Roche Complete EDTA-free tablets]). Membranes were then extracted from the homogenates by ultracentrifugation. The extracted crude membrane pellet was collected, resuspended in 1×PBS to a final concentration of 5 mg wet membrane per ml 1×PBS and crosslinked with 5 mM BS3 (Proteochem), followed by solubilization with a solubilization buffer containing 100 mM (NH_4_)_2_SO_4_, 20 mM Tris-HCl (pH 8.0), 300 mM NaCl, 20 mM imidazole, 1% (wt/vol) Cymal-5 (Anatrace), and a cocktail of protease inhibitors (Roche Complete EDTA-free tablets). The suspension was ultracentrifuged for 30 min at 100,000 × g and 4°C. The supernatant was collected and was mixed with a small volume of preequilibrated Ni-nitrilotriacetic acid (NTA) beads (Qiagen) for 2 h on a rocking platform at 4°C. The mixture was then injected into a small column and washed with a buffer containing 20 mM Tris-HCl (pH 8.0), 100 mM (NH_4_)_2_SO_4_, 1 M NaCl, 30 mM imidazole, and 0.5% Cymal-5. The beads were resuspended in a buffer containing 20 mM Tris-HCl (pH 8.0), 100 mM (NH_4_)_2_SO_4_, 250 mM NaCl, 4.5 mg/ml Amphipol A8-35 (Anatrace), 0.006% DMNG (Anatrace) and a cocktail of protease inhibitors (Roche Complete EDTA-free tablets), and incubated for 2 hours on a rocking platform. The mixture was applied to a column and the buffer was allowed to flow through. The beads were then resuspended in a buffer containing 20 mM Tris-HCl (pH 8.0), 100 mM (NH_4_)_2_SO_4_, 250 mM NaCl, 4.5 mg/ml Amphipol A8-35 (Anatrace) and a cocktail of protease inhibitors (Roche Complete EDTA-free tablets), and incubated for 2 hours on a rocking platform. The mixture was added to a column and the buffer allowed to flow through, followed by washing with 10 bed volumes of a buffer containing 20 mM Tris-HCl (pH 8.0), 100 mM (NH_4_)_2_SO_4_, and 250 mM NaCl. Proteins were eluted from the bead-filled column with a buffer containing 20 mM Tris-HCl (pH 8.0), 100 mM (NH_4_)_2_SO_4_, 250 mM NaCl, and 250 mM imidazole. The buffer of the eluted Env(-) glycoprotein solution was exchanged with imaging buffer containing 20 mM Tris-HCl (pH 8.0), 100 mM (NH_4_)_2_SO_4_, and 250 mM NaCl with a Centrifugal Filter (Millipore), and was concentrated. Before cryo-plunging, Cymal-6 (Anatrace) was added to the Env(-) glycoprotein solution at a final concentration of 0.005%.

### Single-molecule FRET: sample preparation, data acquisition and analysis

Analysis of the conformational dynamics of HIV-1 Env was performed after enzymatic labeling of the V1 and V4 regions of gp120 on the purified (His)_6_-tagged HIV-1_JR-FL_ Env(-) glycoprotein with Cy3 and Cy5 fluorophores, respectively, as previously described^5^. A transfection ratio of 20:1 of non-tagged: V1/V4-tagged HIV-1_JR-FL_ Env(-) was used to ensure that only one protomer within a trimer carries enzymatic tags for site-specific labeling. The purified HIV-1_JR-FL_ Env(-) glycoprotein in buffer (20 mM Tris-HCl (pH 8.0), 10 mM MgCl_2_, 10 mM CaCl_2_, 100 mM (NH_4_)_2_SO_4_, 250 mM NaCl, 0.005% Cymal-6, 10 μM BMS-806) was labeled with Cy3B(3S)-cadaverine (0.5 μM), transglutaminase (0.65 μM; Sigma Aldrich), LD650-CoA (0.5 μM) (Lumidyne Technologies), and AcpS (5 μM) at room temperature overnight. After labeling, Env(-) trimers were purified using Zeba^TM^ spin desalting columns (Thermo Fisher) to remove free dyes. Finally, prior to imaging, fluorescence-labeled HIV-1_JR-FL_ Env(-) carrying the (His)_6_ epitope tag was incubated with biotin-conjugated anti-(His)_6_ tag antibody (HIS.H8, Invitrogen) at 4° for two hours.

All smFRET data were acquired on a home-built total internal reflection fluorescence (TIRF) microscope, as previously described.^5, 55^ Fluorescently labeled HIV-1_JR-FL_ Env(-) trimers were immobilized on passivated streptavidin-coated quartz microscopy slides and washed with pre-imaging buffer specifically made for this experiment. The pre-imaging buffer consisted of 20 mM Tris-HCl (8.0), 100 mM (NH_4_)_2_SO_4_, 250 mM NaCl, 0.005% Cymal-6, and 10 μM BMS-806. For smFRET analysis, a cocktail of triplet-state quenchers and 2 mM protocatechuic acid (PCA) with 8 nM protocatechuic 3,4-dioxygenase (PCD) was added to the above pre-imaging buffer to remove molecular oxygen. Cy3 and Cy5 fluorescence was detected with a 60x water-immersion objective (Nikon), split by a diachronic mirror (Chroma), and imaged on two synchronized ORCA-Flash4.0v2 sCMOS cameras (Hamamatsu) at 40 frames/second for 80 seconds.

smFRET data analysis was performed on the customized Matlab (Mathworks) program SPARTAN.^55^ Fluorescence intensity trajectories were extracted from recorded movies, and FRET efficiency (FRET) was calculated based on FRET= I_A_/(γI_D_+I_A_), where I_D_ and I_A_ are the fluorescence intensities of donor (D) and acceptor (A), respectively, and γ is the correlation coefficient, which incorporates the difference in quantum yields of donor and acceptor and detection efficiencies of the donor and acceptor channels. FRET trajectories were further compiled into a FRET histogram, which provides information about the distribution of Env(-) molecules among the conformational states. The state distributions in the FRET histogram were then fitted to the sum of three Gaussian distributions (based on previously identified FRET trajectories) in Matlab, and the occupancy of each state was further obtained from the area under each Gaussian distribution.

### Immunoprecipitation of cell-surface Env

One day prior to transfection, HOS cells were seeded in 6-well plates (6 × 10^5^ cells/well). The cells were transfected the next day with 0.4 µg of the pSVIIIenv plasmid expressing the wild-type HIV-1_JR-FL_ Env and 0.05 µg of a Tat-expressing plasmid. Two days later, the cells were washed twice with blocking buffer (1×PBS with 5% FBS) and then incubated for 1 hour at 4 °C with 6 μg/μl anti-gp120 monoclonal antibody. Cells were then washed four times with blocking buffer, four times with washing buffer (140 mM NaCl, 1.8 mM CaCl_2_, 1 mM MgCl_2_ and 20 mM Tris, pH 7.5), and lysed in NP-40 buffer (0.5 % NP-40, 0.5 M NaCl and 10 mM Tris, pH 7.5) for 5 min at 4°C with gentle agitation. Lysates were cleared by centrifugation at 15,000xg for 30 min at 4°C. Antibody-bound Env was precipitated using Protein A-Sepharose beads and analyzed by SDS-PAGE and Western blotting with a horseradish peroxidase (HRP)-conjugated rabbit anti-gp120 polyclonal serum.

### Cell-based enzyme-linked immunosorbent assay (ELISA)

CHO cells expressing HIV-1_JR-FL_ Env(-) were induced with 1 μg/ml doxycycline with or without 10 μM BMS-806. Fifteen to twenty-four hours later, the cells were washed twice with washing buffer #1 (20 mM Hepes, pH 7.5, 1.8 mM CaCl_2_, 1 mM MgCl_2_, 140 mM NaCl), and crosslinked with 5 mM BS3 or incubated in buffer without crosslinker. Forty-five minutes later, the cells were quenched with quench buffer (50 mM Tris, pH 8.0, 1.8 mM CaCl_2_, 1 mM MgCl_2_, 140 mM NaCl). The cells were blocked with a blocking buffer (35 mg/ml BSA, 10 mg/ml non-fat dry milk, 1.8 mM CaCl_2_, 1 mM MgCl_2_, 25 mM Tris, pH 7.5 and 140 mM NaCl) and incubated with the indicated primary antibody in blocking buffer for 30 min at 37°C. Cells were then washed three times with blocking buffer and three times with washing buffer #2 (140 mM NaCl, 1.8 mM CaCl_2_, 1 mM MgCl_2_ and 20 mM Tris, pH 7.5) and re-blocked with the blocking buffer. A horseradish peroxidase (HRP)-conjugated antibody specific for the Fc region of human IgG was then incubated with the samples for 45 min at room temperature. Cells were washed three times with blocking buffer and three times with washing buffer #2. HRP enzyme activity was determined after addition of 35 µl per well of a 1:1 mix of Western Lightning oxidizing and luminal reagents (Perkin Elmer Life Sciences) supplemented with 150 mM NaCl. Light emission was measured with a Mithras LB940 luminometer (Berthold Technologies).

### Analysis of Env(-) glycoforms in BMS-806-treated cells

CHO cells expressing HIV-1_JR-FL_ Env(-) were treated with 1 µM BMS-806 or an equivalent volume of the carrier, DMSO. After 18-24 h of culture, the cells were harvested and lysed in homogenization buffer (see above) and treated with different glycosidases following the manufacturer’s instructions. The lysates were analyzed by Western blotting with a horseradish peroxidase (HRP)-conjugated anti-HIV-1 gp120 antibody, as described above.

### Analysis of Env glycopeptides

The sample preparation and mass spectrometric analysis of Env(-) glycopeptides has been described previously^49^, and no changes were made to the procedure for the current analysis. Briefly, the Env(-) glycoprotein was denatured with urea, reduced with TCEP, alkylated with iodoacetamide, and quenched with dithiothreitol. The protein was then buffer exchanged, digested with trypsin alone or with a combination of trypsin and chymotrypsin, generating glycopeptides.

The glycopeptides were analyzed by LC-MS on an LTQ-Orbitrap Velos Pro (Thermo Scientific) mass spectrometer equipped with ETD (electron transfer dissociation) that was coupled to an Acquity Ultra Performance Liquid Chromatography (UPLC) system (Waters). About 35 micromoles of digest was separated by reverse phase HPLC using a multistep gradient, on a C18 PepMap™ 300 column. The mass spectrometric analysis was performed using data-dependent scanning, alternating a high-resolution scan (30,000 at m/z 400), followed by ETD and collision-induced dissociation (CID) data of the five most intense ions. The glycopeptides were identified in the raw data files using a combination of freely available glycopeptide analysis software and expert identification, as described previously^49^.

### Cryo-EM sample preparation and data collection

A 3-μl drop of 0.3 mg/ml Env(-) protein solution was applied to a glow-discharged C-flat grid (R1/1 and R1.2/1.3, 400 Mesh, Protochips, CA, USA), blotted for 2 sec, then plunged into liquid ethane and flash-frozen using an FEI Vitrobot Mark IV. The cryo-grid was imaged in an FEI Tecnai Arctica microscope, equipped with an Autoloader, at a nominal magnification of 21,000 times and an acceleration voltage of 200 kV. Coma-free alignment was manually conducted prior to data collection. Cryo-EM data were collected semi-automatically by Leginon^56^ version 3.1 on the Gatan K2 Summit direct detector camera (Gatan Inc., CA, USA) in a super-resolution counting mode, with a dose rate of 8 electrons/pixel/second and an accumulated dose of 50 electrons/Å^2^ over 38 frames per movie. The calibrated physical pixel size and the super-resolution pixel size are 1.52 Å and 0.76 Å, respectively. The defocus for data collection was set in the range of −1.0 to −3.0 μm. A total of 12,440 movies were collected, from which 10,299 movies were selected for further data analysis after screening and inspection of data quality.

### Data analysis and cryo-EM refinement

The raw movie frames were first corrected for their gain reference and each movie was used to generate a micrograph that was corrected for sample movement and drift with the MotionCor2 program^57^. These drift-corrected micrographs were used for the determination of the actual defocus of each micrograph with the CTFFind4 program^58^. Using DeepEM, a deep learning-based particle extraction program that we recently developed^42^, 1,436,424 particles of Env(-) were automatically selected in a template-free fashion. Reference-free 2D classification was done in a recently developed program, ROME^43^, which combines maximum likelihood-based image alignment^44^ and statistical manifold learning-based classification^43^. 3D classification was conducted with the maximum-likelihood approach implemented in either RELION 1.3 or ROME 1.1.

All 2D and 3D classifications were done at a pixel size of 1.52 A□. After the first round of reference-free 2D classification, bad particles were rejected upon inspection of class average quality, which left 1,366,095 particles. The initial model, low-pass filtered to 60 A□, was used as the input reference to conduct unsupervised 3D classification into 5 classes with *C*3 symmetry, using an angular sampling of 7.5° and a regularization parameter T of 4.

One class, consisting of 49.5% of the particles, gave rise to an averaged reconstruction overall resembling that of antibody Fab-bound sgp140 SOSIP.664 trimer structures. This class was further divided into 6 classes by 3D classification, which were grouped into 3 sub-datasets. Each sub-dataset was further classified into 6 classes. We then obtained 15 good classes and 3 junk classes. The classes showing complete assemblies and conformations generally resembling published Env trimer structures were deemed “good classes”, whereas the classes showing incomplete assemblies, strong loss of three-fold symmetry or random features resulting from noise or misalignment were deemed “junk classes”. Auto-refinement of these good classes in RELION 1.3 with imposition of *C*3 symmetry was performed with a pixel size of 1.52 A□ Twelve classes exhibited a topology similar to that of sgp140 BG505 SOSIP.664 and were designated State P2. Three classes exhibited a topology similar to that of the electron tomographic map observed on native HIV-1 virus particles and were designated State P1. The 12 State P2 classes were combined and further classified into 6 classes without assuming any symmetry. Four classes that exhibited better *C*3 symmetry were further refined with *C*1 symmetry. Two classes containing a total of 121,979 particles showed nearly the same conformation and reached a resolution of 5.5 A□ (gold-standard FSC at 0.143 cutoff) after refinement with imposition of *C*3 symmetry.

The four other initial classes, which together contain 50.5% of the particles, were reclassified into 4 classes with *C*3 symmetry imposed. Two of these classes showed features corresponding to State P1. We therefore combined all classes resembling State P1 and classified them into 4 classes without imposing *C*3 symmetry. Two classes were chosen for further refinement without imposing *C*3 symmetry in RELION 1.3; after refinement, only one class consisting of 11,667 particles showed good *C*3 symmetry. The final reconstruction was done with this class with imposition of *C*3 symmetry, and the resolution was estimated to be 8.0 A☐ by gold-standard FSC at 0.143 cutoff.

### Tilt-pair validation

A set of 1,310 pairs of movies was collected with a Tecnai Arctica transmission electron microscope using the Gatan K2 Summit direct electron detector in a super-resolution counting video mode with the MSI-RCT (Random Conical Tilt) application in Leginon 3.1. A relatively large set of tilt-pair micrographs was collected to ensure that there would be enough particles to enable 3D classification prior to tilt-pair validation. The same sample areas were exposed twice at a pair of tilt angles (−11° and +11°), with a dose of 30 electrons/Å^2^ per exposure. The tilt-pair movies were corrected for drift and local movement with the MotionCor2 program^557^. From the micrograph set taken at the first tilt angle, 100,000 single-particle images of the Env(-)glycoprotein were automatically picked in a template-free fashion using the program DeepEM^42^. The local defocus of each particle was calculated with the program Gctf^59^. 2D and 3D classifications of this particle dataset were processed in a fashion similar to that described above for the particles without tilting. After the first round of reference-free 2D classification, bad classes of particles were rejected upon inspection of the class average quality. The previous initial model, either the State-P1 or State-P2 reconstruction, low-pass filtered to 60 Å, was used as the input reference to conduct an unsupervised 3D classification into 6 classes with *C*3 symmetry imposed, using an angular sampling of 7.5° and a regularization parameter T of 4. Among the 6 output classes, Class 2 and Class 3 reconstructions, consisting of 49.8% and 8.49% of the total particles, exhibited structural features virtually identical to those of State P2 and State P1, respectively. Both classes were obtained exclusively from the tilt-pair data, independent of the data of untilted samples used to obtain the initial P1 and P2 maps previously.

A total of 135 pairs of particles from Class 3, corresponding to the State-P1 conformation, were used for tilt-pair validation analysis. The particle image of the second tilt in each particle pair was picked and verified manually with EMAN 2.2. The dataset of tilt-pair particles was used to perform tilt-pair validation against the previously refined P1 and P2 maps using the e2tiltvalidate.py code from the EMAN package, in which projection matching and tilt distance were calculated with *C*3 symmetry imposed. The results were plotted with a custom-made Python script, while the statistics were computed with the EMValidationPlot function from EMAN.

### Model building and structural analysis

To build the pseudo-atomic model of State P2, we used a previously published PGT151-bound Env(+)ΔCT structure^23^ and then manually improved the main-chain and side-chain fitting in Coot^60^ to generate the starting coordinate files. The fitting of the pseudo-atomic model into the State-P2 density map was further improved through the program Phenix.real_space_refine^61^, with secondary-structure and geometry restraints to prevent overfitting. The pseudo-atomic model of the gp120 subunit from State P2 was used as a whole to perform rigid-body fitting into the State-P1 density. Structural comparison was conducted in Pymol^62^ and Chimera. Interaction analysis between adjacent subunits was performed using PISA^63^. All figures of the structures were produced in Chimera and Pymol^64^.

## Accession numbers

The cryo-EM reconstructions of states P1 and P2 reported in this paper have been deposited in the Electron Microscopy Data Bank under accession numbers EMD-7524 and EMD-7525, respectively. The pseudo-atomic models of P1 and P2 have been deposited in the Protein Data Bank under ID codes 6CMT and 6CMU. The cryo-EM raw data, including the motion-corrected micrographs, the particle stacks of P1 and P2 used for final refinement, the motion-corrected micrographs used for tilt-pair validation and the tilt-pair particle stacks for P1 validation, have been deposited into the Electron Microscopy Pilot Image Archive (www.ebi.ad.uk/emdb/ampiar) under accession no. EMPIAR-10163.

## Author contributions

J.S. and Y.M. conceived this study. H. Ding and J.C.K. prepared the Env(-)-expressing CHO cells. S.Z. and R.T.S. analyzed Env(-) antigenicity and established a purification scheme for the Env(-) protein. S.Z. and W.L.W. screened the samples for optimization of cryo-EM imaging. W.L.W. conducted cryo-electron microscopy, collected all data and preprocessed the data. S.C. performed data analysis and refined the maps. W.L.W. and S.C. performed validation tests. S.Z., S.C. and Y.M. built the pseudo-atomic models. E.P.G., S.Z. and H. Desaire analyzed the Env(-) glycans. M.L. and S.Z. conducted smFRET experiments. Y.M. and J.S. wrote the manuscript. All authors contributed to data analysis and manuscript preparation.

## Acknowledgments

We thank Dr. Walther Mothes (Yale University) for helpful advice. This work was funded in part by NIH grants AI125093 (H. Desaire), AI93256, AI100645 and AI124982 (J.S.), by an Intel academic grant (Y.M.), by a grant of the Thousand Talents Plan of China (Y.M.), by grants from the National Natural Science Foundation of China 11774012 and 91530321 (Y.M.), and by a gift from William F. McCarty-Cooper. The cryo-EM experiments were performed in part at the Center for Nanoscale Systems at Harvard University, a member of the National Nanotechnology Coordinated Infrastructure Network (NNCI), which is supported by the National Science Foundation under NSF award no. 1541959. The cryo-EM facility was funded through the NIH grant AI100645, Center for HIV/AIDS Vaccine Immunology and Immunogen Design (CHAVI-ID). The data processing was performed in part in the Sullivan cluster, which is supported by a gift from Mr. and Mrs. Daniel J. Sullivan, Jr.

