## Supplementary Materials for "Structural insights into the conformational plasticity of the full-length trimeric HIV-1 envelope glycoprotein precursor"

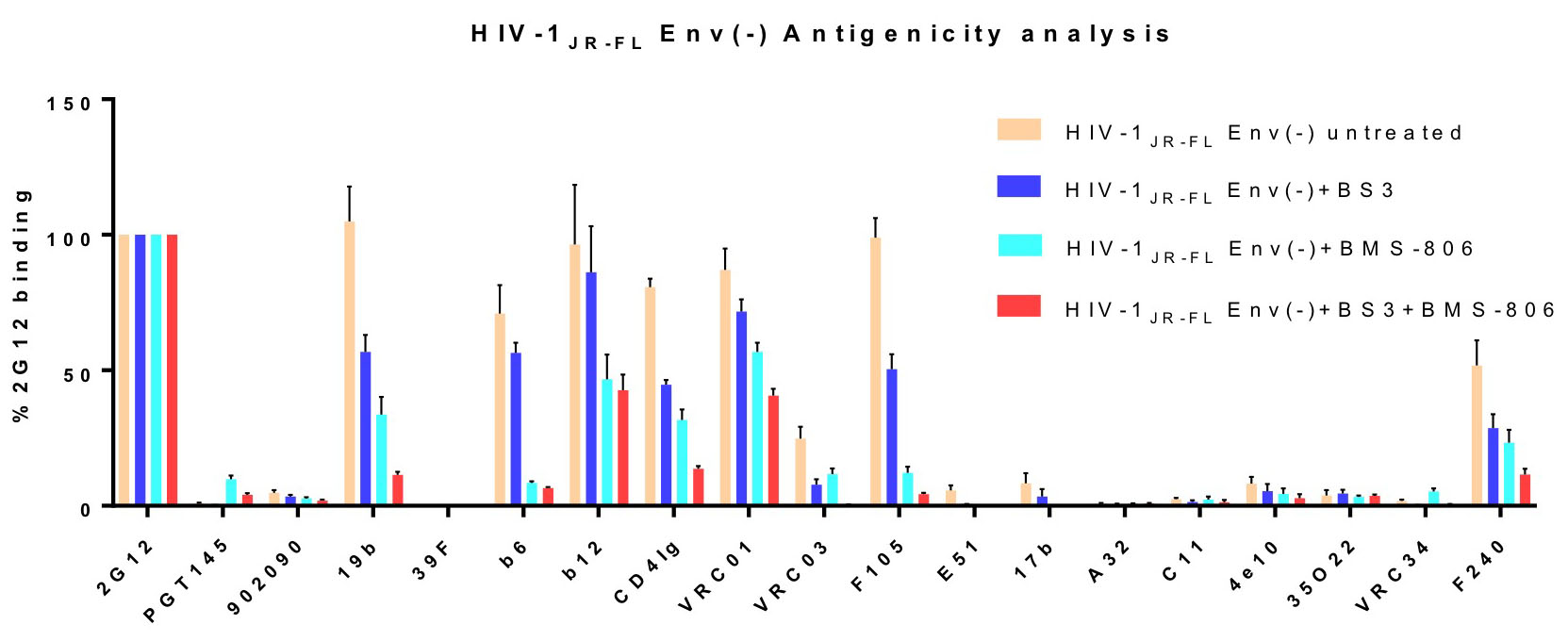

**Supplementary Figure 1**. **Antigenic profile of full-length HIV-1_JR-FL_ Env(-) expressed on the surface of CHO cells.** The effect of BS3 or/and BMS-806 treatment on HIV-1_JR-FL_ Env(-) antigenicity was evaluated by cell-based ELISA. All values were normalized against 2G12 binding and derived from at least three independent experiments. Note that the HIV-1_JR-FL_ Env(-) glycoprotein is not recognized by the PGT145 antibody, which serves as a negative control.

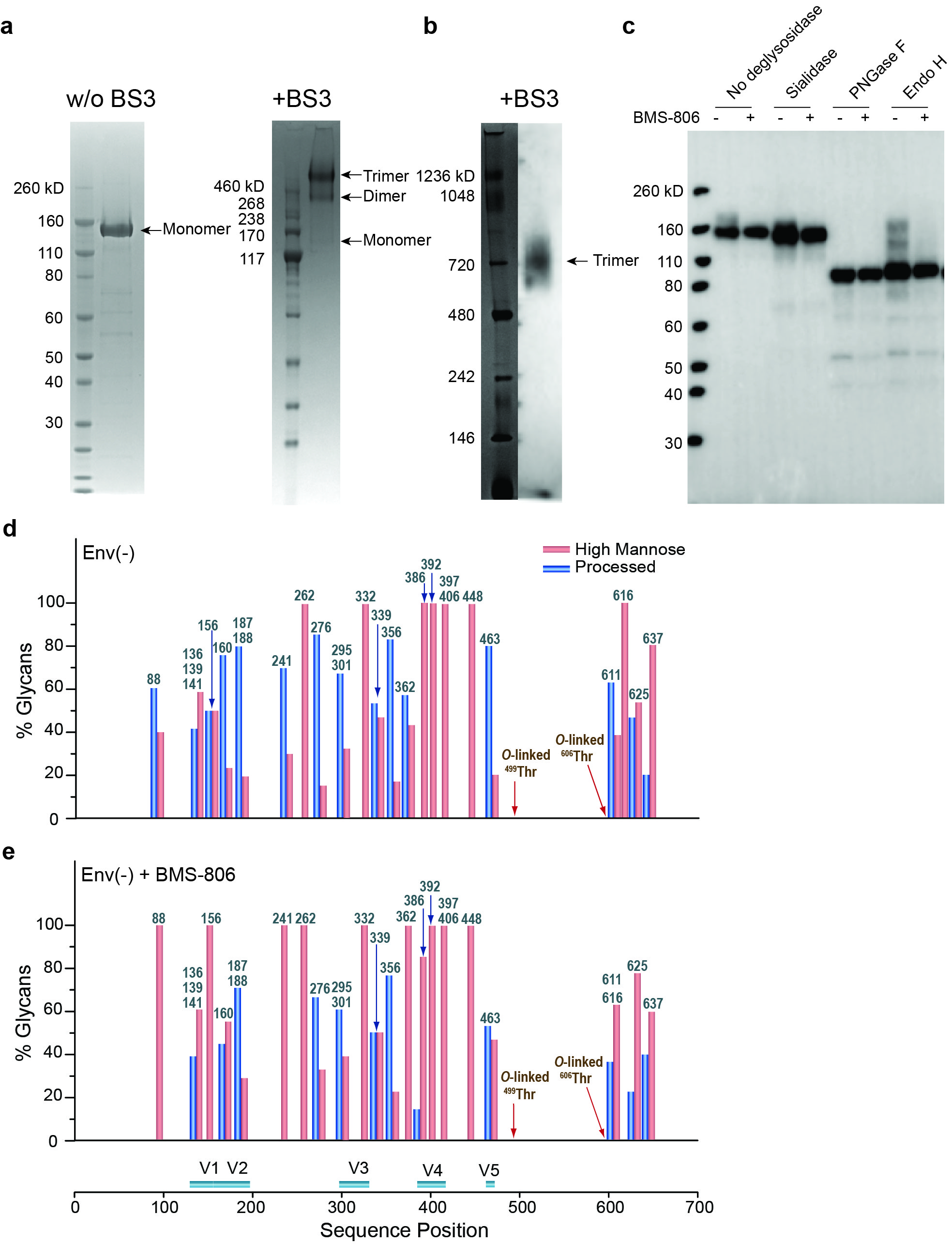

**Supplementary Figure 2**. **Characterization of the full-length Env(-) in cell lysates and in detergent-solubilized purified forms.** (**a**) Purified HIV-1_JR-FL_ Env(-) without and with crosslinking by BS3 was run on a NUPAGE 4-12% BT gel stained by Coomassie Blue. (**b**) Purified HIV-1_JR-FL_ Env(-) crosslinked by BS3 was run on a NativePAGE 4-16%BT gel and subjected to Western blotting with an HRP-conjugated anti-HIV-1 gp120 antibody. (**c**) The effect of BMS-806 on HIV-1_JR-FL_ Env(-) glycosylation was evaluated by Western blotting after deglycosylase digestion. The purified HIV-1_JR-FL_ Env(-) glycoproteins were digested with the indicated deglycosylases, run on a NUPAGE 4-12% BT gel, and subjected to Western blotting with an HRP-conjugated anti-HIV-1 gp120 antibody. The results shown are representative of those obtained in three independent experiments. Note that BMS-806 treatment decreases Env(-) heterogeneity by reducing the levels of Endo H-resistant glycoforms. (**d**, **e**) The bar graphs show the glycan profiles at each glycosylation site of HIV-1_JR-FL_ Env(-) purified from untreated CHO cells (**d**) or CHO cells treated with 10 µM BMS-806 (**e**), as determined by mass spectrometry. The glycan composition (in percent) was broadly characterized as high-mannose (red bars) or processed glycans (blue bars).

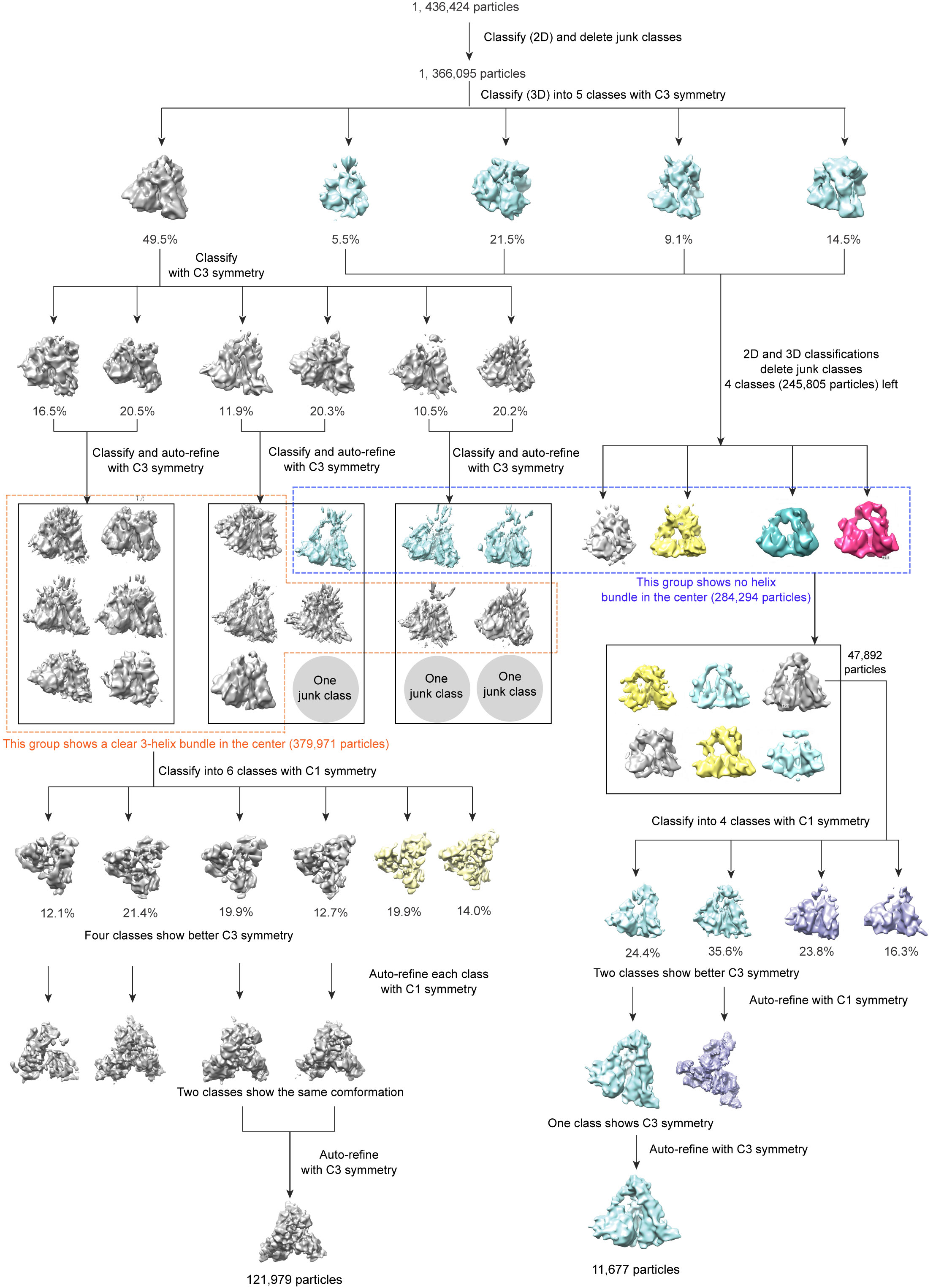

**Supplementary Figure 3**. **Multiple rounds of 3D classification and refinement leading to the reconstructions of the Env(-) P1 and P2 states.** One round of rough 3D classification and a round of refined classification followed by auto-refine separated the particles into two groups, both with C3 imposed. Recombination of the class groups was based on the emerged map characteristics including the central cavity and three-helix bundle. The two groups were further classified and auto-refined with both C1 and C3 symmetry to remove heterogeneous particles that are potentially damaged or in a state deviating substantially from the most self-consistent conformations.

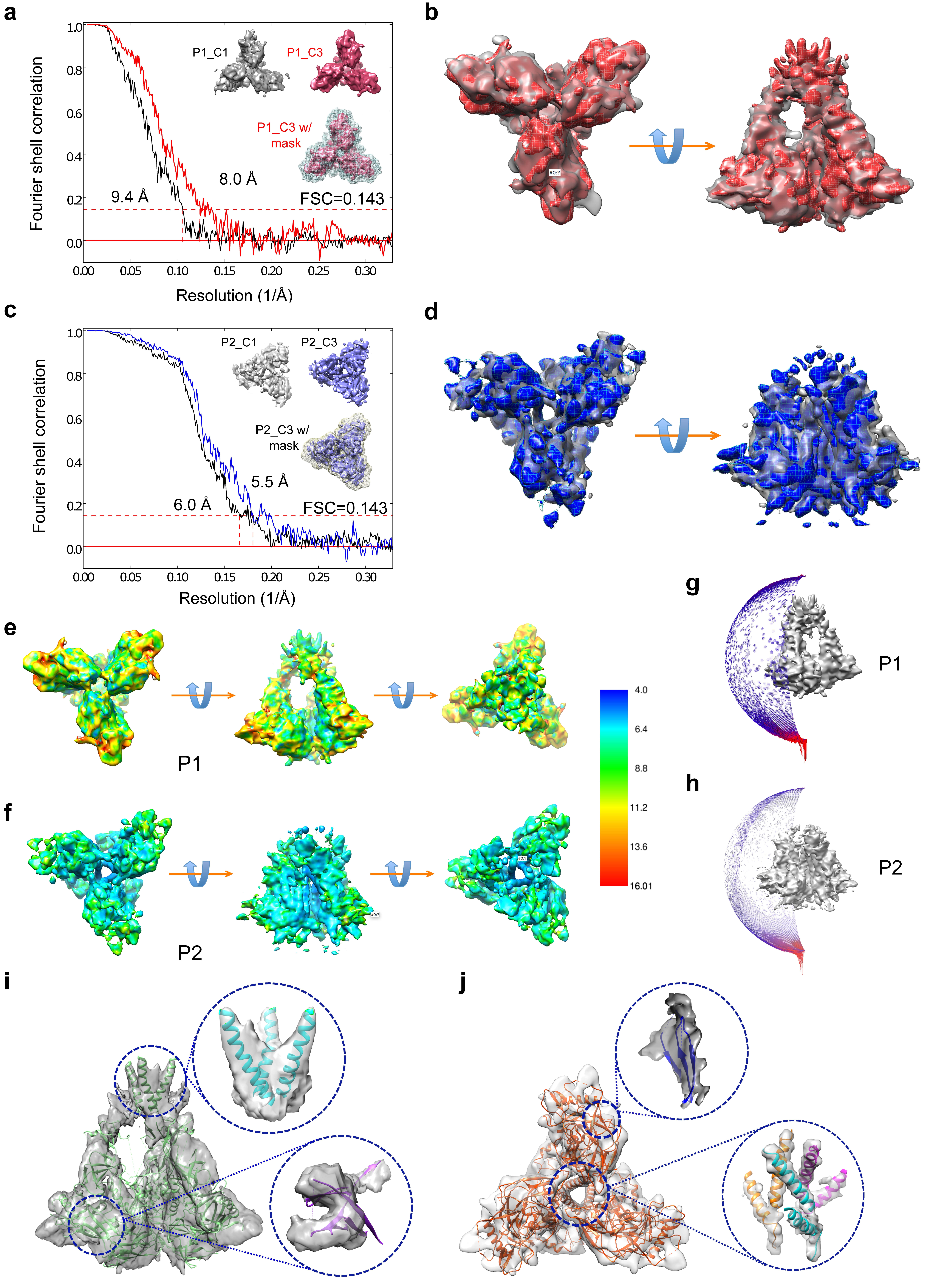

**Supplementary Figure 4**. **Cryo-EM structural determination of the Env(-) P1 and P2 states.** **(a)** Comparison of the final maps of the Env(-) P1 state obtained without imposing C3 symmetry (gray) and with C3 symmetry imposed (red). Resolutions were estimated by gold-standard Fourier shell correlation (FSC 0.143). The 3D mask used for the FSC measurement is shown as a grey mesh in the lower insert. **(b)** The superposed C1 (gray) and C3 (red) State-P1 density maps are shown from the perspective of the target cell (left) and from the side (right). (**c**) Comparison the final maps of the Env(-) P2 state without imposing C3 symmetry (gray) and with C3 symmetry imposed (blue). The 3D mask used for the FSC measurement is shown as a grey mesh in the lower insert. (**d**) The superposed C1 (gray) and C3 (blue) State-P2 density maps are shown from the perspective of the target cell (left) and from the side (right). (**e**) and (**f**) show the local resolution of the P1 and P2 Env(-) maps, respectively. The maps are colored according to the local resolution, indicated by the color gradient (units in Angstroms). (**g**) and (**h**) 3D reconstructions of the P1 and P2 Env(-) states, respectively, and the orientation distributions of the aligned particles used for the reconstructions. The particles exhibited a preferred orientation on the cryo-grids, with the trimer axis approximately parallel to the z-axis. The addition of Cymal-6 to the purified Env(-) glycoprotein prior to cryo-plunging diminished but did not completely eliminate this orientation preference. (**i**) and (**j**) The unsharpened maps of the P1 and P2 Env(-) are superimposed with their respective pseudo-atomic models.

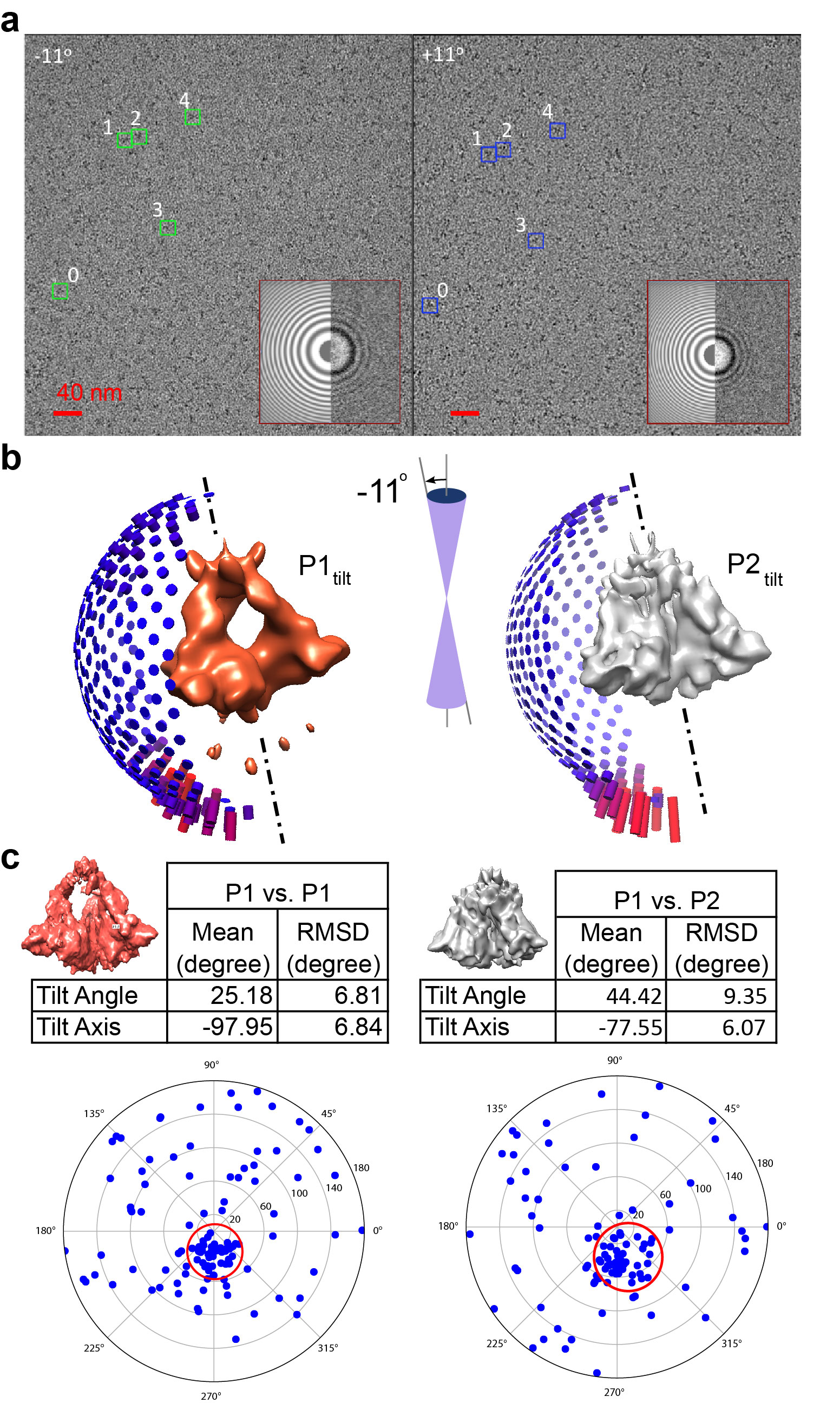

**Supplementary Figure 5**. **Tilt-pair validation of the Env(-) State-P1 map.** (**a**) Representative micrographs of the tilt-pair data set, from which five pairs of P1 particles were picked (labeled boxes). The stage tilt of the micrograph on the left was -11°, and the stage tilt of the micrograph on the right was +11°. Insets show comparisons of the 2D Fourier transforms calculated from the experimental data (right half circles) and the fitted contrast transfer functions (left half circles). (**b**) P1 and P2 particles were identified by 3D classification in micrographs of the first tilt (-11^o^), such as that shown in (**a**). The resultant maps of P1 and P2 are shown here with the orientation distribution of the corresponding particles. (**c**) Statistics and plots of the tilt-pair validation of the State-P1 map. The P1 particle pairs were aligned against the P1 model (left) or the P2 model (right). Both models used were those obtained from the un-tilted micrographs. The position of each blue dot represents the direction and the amount of tilt (radial direction) for a particle pair in polar coordinates. Both results show a clustering of dots in the direction of the stage tilt; however, the use of the P1 reference map (left) resulted in tighter clustering and a mean tilt angle of 25.18, closer to the known stage tilt angle of 22°. By contrast, a tilt angle of 44° was predicted when the P1 particle set was calculated against the P2 map (right). The results on the left validate the P1 model and confirm its handedness. The comparison with the results on the right shows that the tilt pair calculation resolves the two related yet distinctive states, P1 and P2.

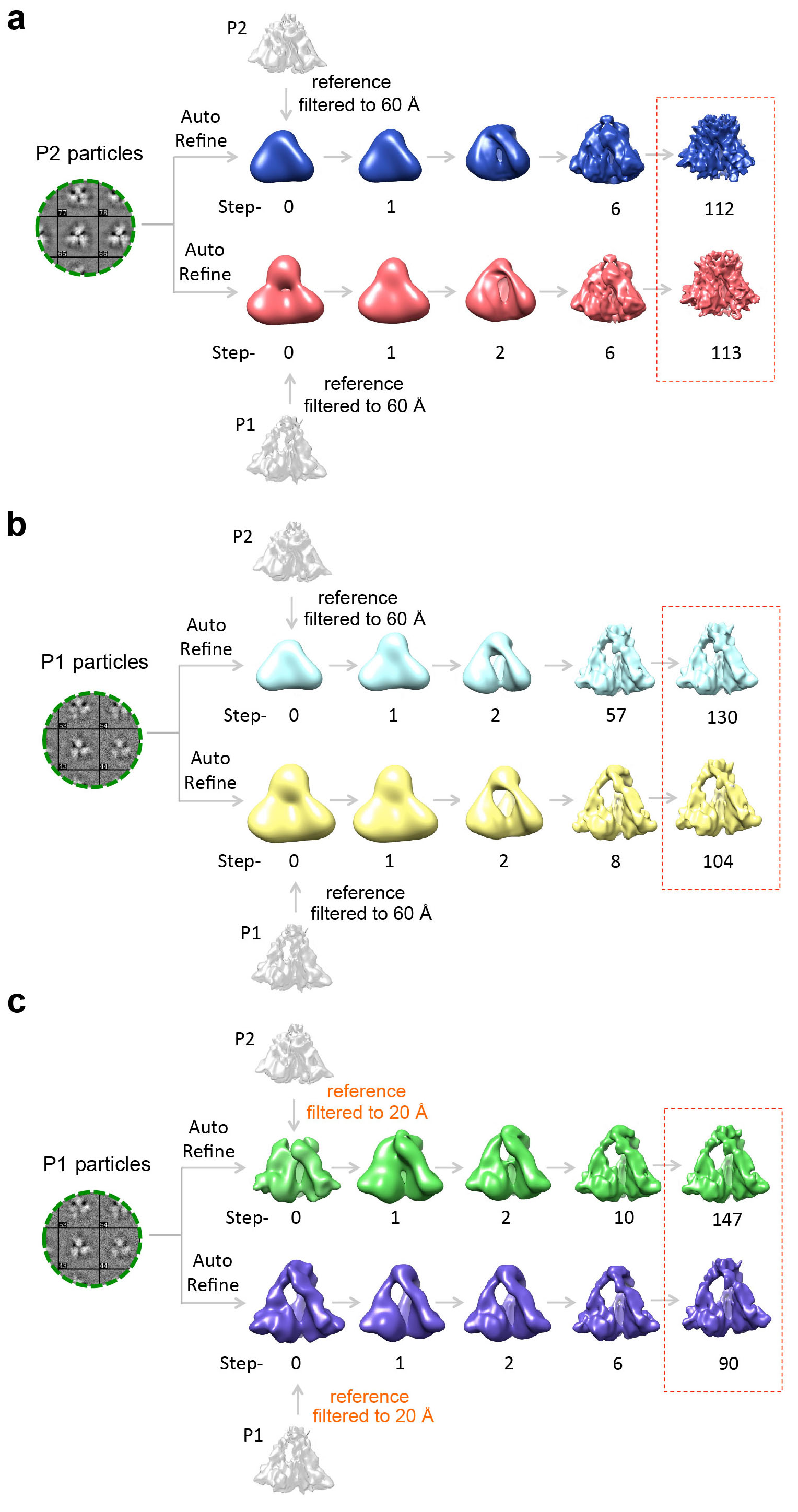

**Supplementary Figure 6**. **Check on the sensitivity of the final Env(-) maps to initial model bias.** (**a**) Comparison of two auto-refine runs of P2 particles with the Env(-) P2 and P1 maps as initial models, both filtered to 60 Å, a commonly used initial model resolution for auto-refine. Both calculations converged to a map corresponding to State P2. This demonstrates that the final map is driven primarily by the input particle data and is relatively insensitive to initial model bias. (**b**) Comparison of two auto-refine runs of P1 particles with the P2 and P1 maps as initial models, both filtered to 60 Å. (**c**) The P1 particles were auto-refined with the Env(-) P2 and P1 maps as initial models, both filtered to 20 Å. The use of a 20-Å initial model introduces more-than-usual initial model bias. Nonetheless, both auto-refine runs still converged to a State-P1 conformation, with P1 characteristics emerging in just a few steps.

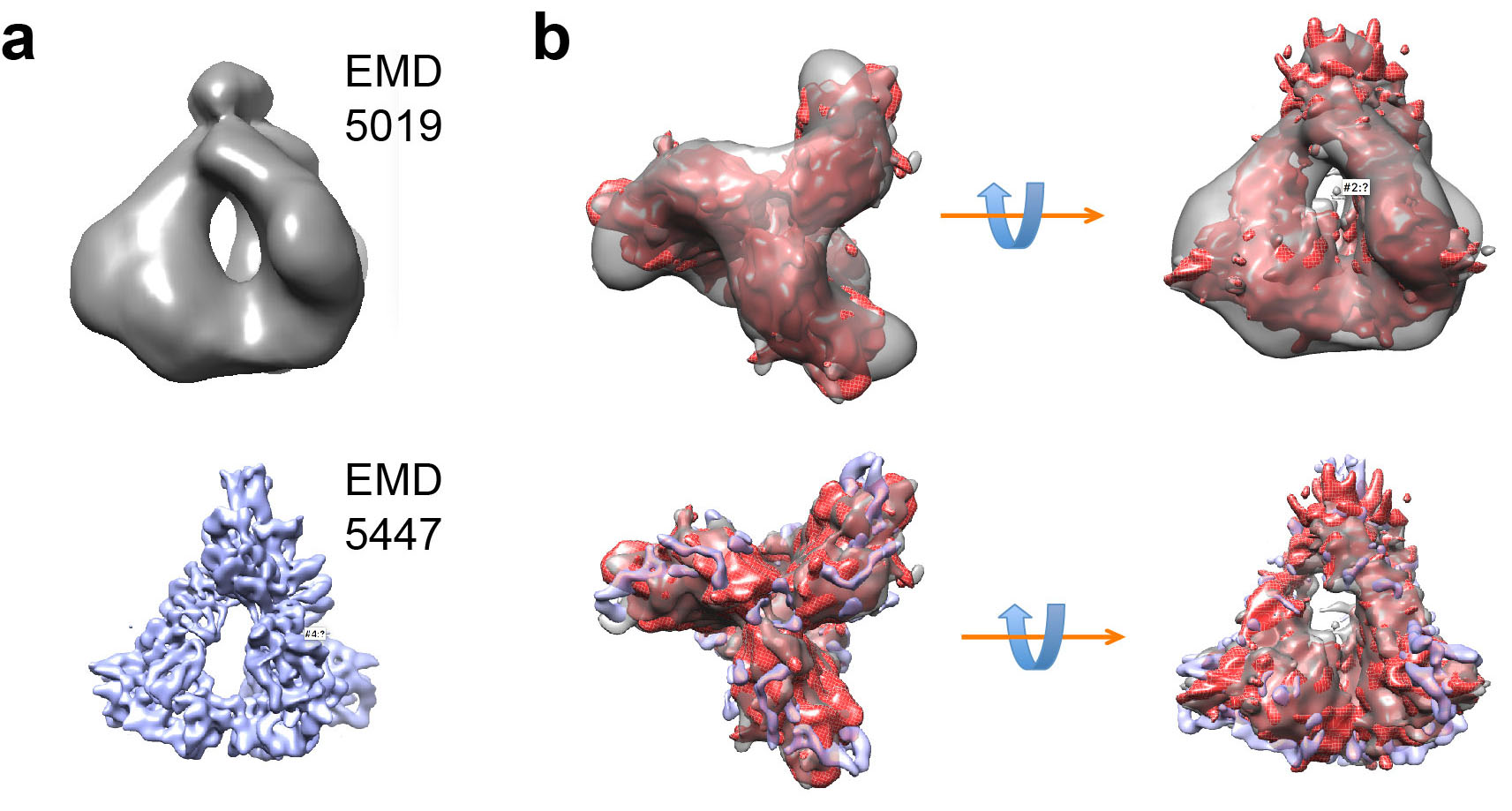

**Supplementary Figure 7**. **Comparison of the State-P1 map to previously published models.** (**a**) An electron tomographic map of the mature HIV-1 Env trimer on virions (EMD-5019) and a cryo-EM map of the Env(-)Δ712 trimer purified from cell membranes (EMD-5447) are shown. (**b**) The State-P1 map (red) is aligned with EMD-5019 (gray) (upper panel) and EMD-5447 (blue) (lower panel), with views from the perspective of the target cell (left) and from the side (right). These models generally conform to each other up to the minimum claimed resolution, and share the central cavity as a common feature.

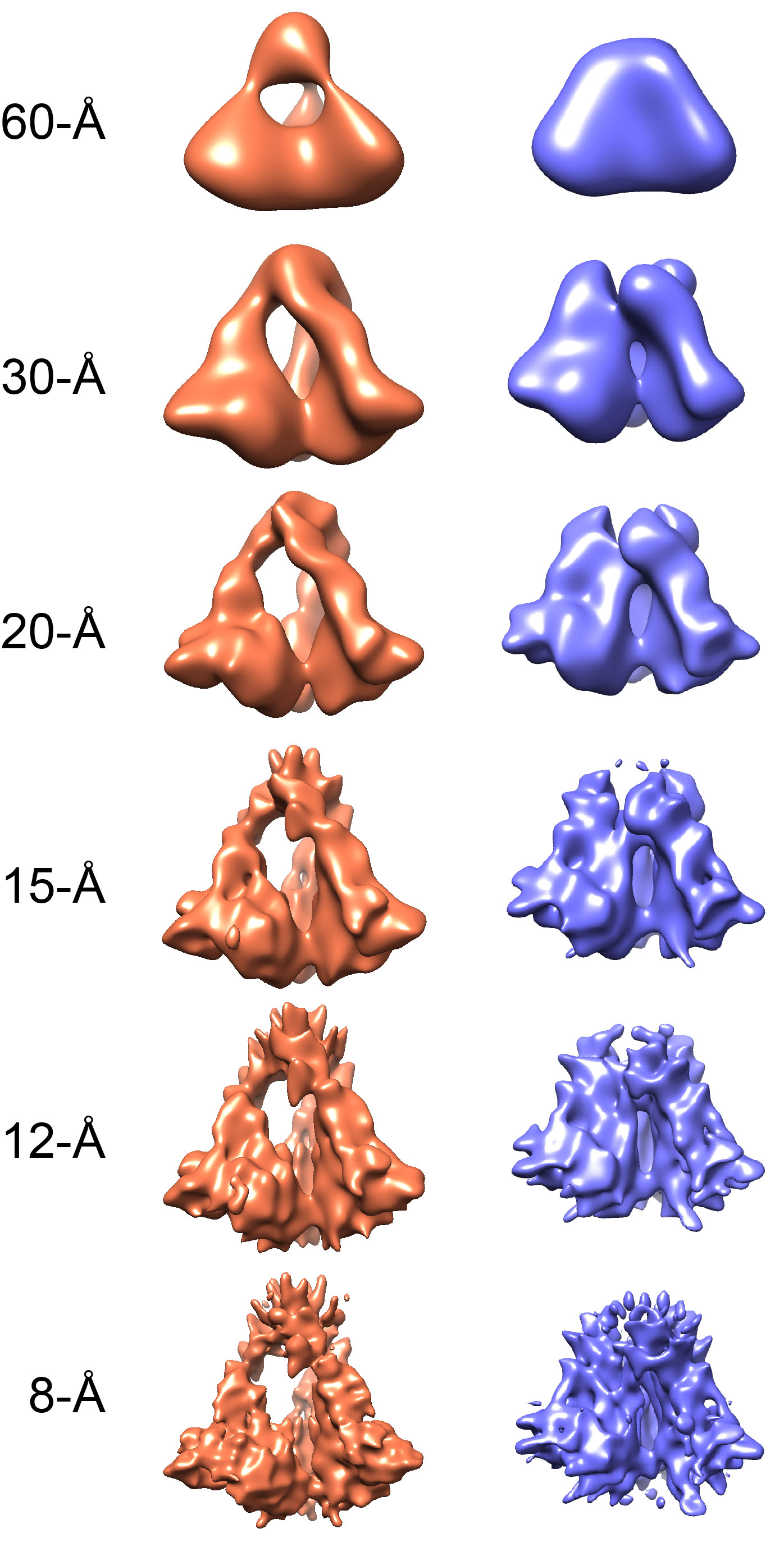

**Supplementary Figure 8. Comparison of the Env(-) State-P1 and State-P2 maps at various filtered resolutions.** The Env(-) State-P1 (orange) and State-P2 (blue) maps were filtered to the indicated resolutions. The central cavity, a main feature of State P1, is present regardless of the chosen filtering resolution and distinguishes State P1 from State P2.

| **Table S1. Statistics of the full-length HIV-1 envelope trimer precursor determined by single-particle cryo-EM** | | |
| --- | --- | --- |
| Electron energy (kV) | 200 | |
| Electron dose (e-/ Å^2^) | 50 | |
| Pixel size corresponding to the physical detector sensor (Å) | 1.52 | |
| Pixel size in the super-resolution counting mode of K2 Summit (Å) | 0.76 | |
| Defocus range (μm) | -3.5 to -1.5 | |
| Number of micrographs | 10,299 | |
| Number of particles | 1,366,095 | |
|  | **P1 state** | **P2 state** |
| Number of particles | 11,677 | 121,979 |
| Resolution (Å) | 8.0 | 5.5 |
| B-factor (Å) | -66 | -70 |
| **Pseudo-crystallographic refinement of atomic models** |  |  |
| Cell dimension a,b,c (Å) |  | 212.8 |
| Cell angle |  | 90 |
| Space group |  | 1 |
| Number of atoms |  | 13320 |
| **Geometric Parameters (RMSD)** |  |  |
| Bond length (Å) | 0.0027 | 0.0031 |
| Bond angle (°) | 0.69 | 0.78 |
| **Ramachandran plot statistics** |  |  |
| Favored (%) | 92.77 | 91.79 |
| Allowed (%) | 5.28 | 6.63 |
| Outliers (%) | 1.95 | 1.59 |
| **MolProbity validation** |  |  |
| Rotamer outliers (%) | 0.23 | 0.22 |
| Clashscore | 12.66 | 15.76 |
| MolProbity score | 2.07 | 2.19 |
| C-beta outliers | 0 | 0 |
